# Total body irradiation primes CD19-directed CAR T cells against large B-cell lymphoma

**DOI:** 10.1101/2025.03.17.643462

**Authors:** Mohammad Alhomoud, Michelle Foley, Mayumi Sugita, Joshua A Fein, Samuel Yamshon, Leandro Martinez, Kai Rejeski, Maider Astorkia, Doron Betel, Renier Brentjens, Koen van Besien, Lorenzo Galluzzi, Olivier Boyer, Jeremie Martinet, Silvia Formenti, Monica L Guzman

## Abstract

CD19-targeting chimeric antigen receptor T cells (CART19) have demonstrated significant effectiveness in treating relapsed or refractory large B-cell lymphoma (LBCL). However, they often fail to sustain durable remissions in more than half of all treated patients. Therefore, there is an urgent need to identify approaches to enhance CART19 efficacy. Here, we studied the impact of low-dose radiation on CART19 activity *in vitro* and find that radiation enhances the cytotoxicity of CART19 against LBCL by upregulating death receptors. Disrupting the FAS receptor diminishes this benefit, indicating that this pathway plays an important role in enhancing the cytotoxic effects of CAR T cells. To further validate these findings, we conducted *in vivo* studies using a lymphoma syngeneic mouse model delivering total body irradiation (TBI). We observed that delivering TBI at a single dose of 1Gy prior to CAR T cell infusion significantly improved CART19-mediated tumor elimination and increased overall survival rates. Importantly, we characterized several important effects of TBI, including enhanced lymphodepletion, improved T cell expansion and persistence, better intra-tumoral migration, and a more favorable, anti-tumor phenotypic composition of the T cells. In summary, for the first time, we have demonstrated preclinically that administering TBI before CART19 infusion significantly accelerates tumor elimination and improves overall survival. This approach holds promise for translation into clinical practice and serves as a valuable foundation for further research to enhance outcomes for patients receiving CART19 treatment.

## INTRODUCTION

CD19 chimeric antigen receptor T cell (CART19) therapy represents a potent treatment strategy for a range of advanced B-cell malignancies, with complete response (CR) rates exceeding 70% for Large B-cell Lymphoma (LBCL) patients in clinical trials [1–3]. However, the challenge lies in achieving durable remission, observed in less than half of LBCL patients [4, 5]. In addition, a subset of patients faces primary refractory disease. This underscores the critical need for a comprehensive understanding of the mechanisms governing response or resistance, in order to facilitate clinical translation of novel CART19 strategies.

Radiotherapy is often utilized to eliminate tumor cells and has also been shown to activate the immune system through several mechanisms. Studies have suggested that tumor-directed radiation primes tumor-reactive CD8^+^ T cells by increasing type I interferon (IFN) induction and enhancing IFN-related gene signatures such as the STING pathway [6–8]. Furthermore, the stress effect of radiation on tumor cells enhances the release of nuclear protein high-mobility group box-1 (HMGB1), boosts adenosine triphosphate (ATP) production, and promotes cell surface translocation of calreticulin (best known as CRT) [9]. These are all essential steps for dendritic cell activation and cross-presentation of tumor-derived antigens to T cells, ultimately resulting in immunogenic cell death [10].

The established effects of radiation therapy on cancer immunotherapy (e.g., immune checkpoint inhibitors) in solid tumors [11–15] have sparked growing interest in its potential impact on CAR T cell therapy. However, the knowledge gap on how radiation specifically influences CAR T cell efficacy remains. In a mouse model of pancreatic adenocarcinoma, it was observed that localized low-dose radiation (LDR) can elicit antigen-independent killing by CAR T cells, ascribed to upregulation of TNF-related apoptosis-inducing ligand (TRAIL) death receptor [16]. We recently described that total body irradiation (TBI) at a single dose of 1Gy enhances the efficacy of CART19 in an immunodeficient humanized mouse model of acute lymphoblastic leukemia (ALL) [17].

For LBCL, the potential of radiation therapy to enhance CART19 therapy has been explored in several retrospective studies [18–27] assessing the impact of radiation when used as a bridging therapy while the patient awaits CART19 manufacturing. Because of the retrospective nature of these investigations, it is difficult to interpret the results in view of the heterogeneity of radiation doses, delivery methods, and timing. Additionally, the absence of correlative studies hampers the interpretation of the impact of radiation on CART19 behavior, warranting preclinical research.

The primary goal of this study was to assess how radiation, in the range of 0.5 and 2Gy influences the *in vitro* cytotoxicity of CART19, and when delivered as total body irradiation (TBI) affects the *in vivo* efficacy of CART19 in a syngeneic B cell lymphoma mouse model. Additionally, we aimed to investigate the role of TBI in enhancing CART19 cell expansion, persistence, intra-tumoral migration, and phenotypic differentiation, as well as its potential influence on tumor cell death signaling pathways that could modulate CART19 activity.

The study findings represent the foundation for a first-in-human prospective phase I/II clinical trial examining the safety and efficacy of TBI (1Gy) in CART19 recipients to treat B cell lymphoma (currently under IRB review, #23-10026672).

## RESULTS

### LDR increases the expression of death receptors (DRs) in LBCL cells and enhances CART19 anti-tumor efficacy *in vitro*

We previously established the ability of radiation (1Gy) to upregulate DRs in ALL cells and potentiate the efficacy of CART19 against irradiated cells *in vitro* and in an immunodeficient mouse model of ALL [17]. Thus, we aimed to determine whether this could be extended to LBCL. To this end, we first assessed the sensitivity of lymphoma cells to radiation by exposing a panel of eight human LBCL cell lines, including both ABC-LBCL (OCI-LY10, SUDHL-2, and RC-K8) and GCB-LBCL (SUDHL-4, SUDHL-5, SUDHL-6, OCI-LY1, and OCI-LY4), as well as a mouse-derived B cell lymphoma cell line (A20 expressing GFP and luciferase; A20 GFP-LUC). Cells were exposed to a single fraction of 1Gy and cell viability was measured by multiparameter flow cytometry after 24 hours. Since we found that 1Gy radiation exposure had minimal effect on the viability of LBCL cells (**Fig. 1A**), we proceeded to assess the expression of CD19 and the DRs, Fas cell surface death receptor (FAS) and TNF receptor superfamily member 10b (TNFRSF10B, best known as DR5 or TRAIL-R2), 24 hours after exposure. We found that a single dose of 1Gy significantly increased the expression of FAS and TRAIL-R2 across all cell lines tested, FAS being the DR with the highest fold change (**Fig. 1B, supplementary Fig. 1A).** Interestingly, in addition to the upregulation of FAS and TRAIL-R2 we found a significant increase in CD19 antigen density in response to 1Gy radiation in LBCL cell lines (**Fig. 1B**), a finding that has been linked to enhanced CART19 efficacy [28]. To better understand the kinetics of FAS and TRAIL-R2 expression over time, we exposed OCI-LY10 cells irradiated with a single dose of 1 Gy and assessed FAS and TRAIL-R2 expression at multiple time points post-radiation. While FAS upregulation started at 3 hours after LDR, peaked at 9 hours, and persisted after 24 hours, TRAIL-R2 began to increase 6 hours post-radiation and remained stable 24 hours later (**supplementary Fig. 1B**). These data suggest that the stability on the effects of radiation on lymphoma cells should allow a 24-hour period to combine with CAR T cells approaches.

**Figure 1.**
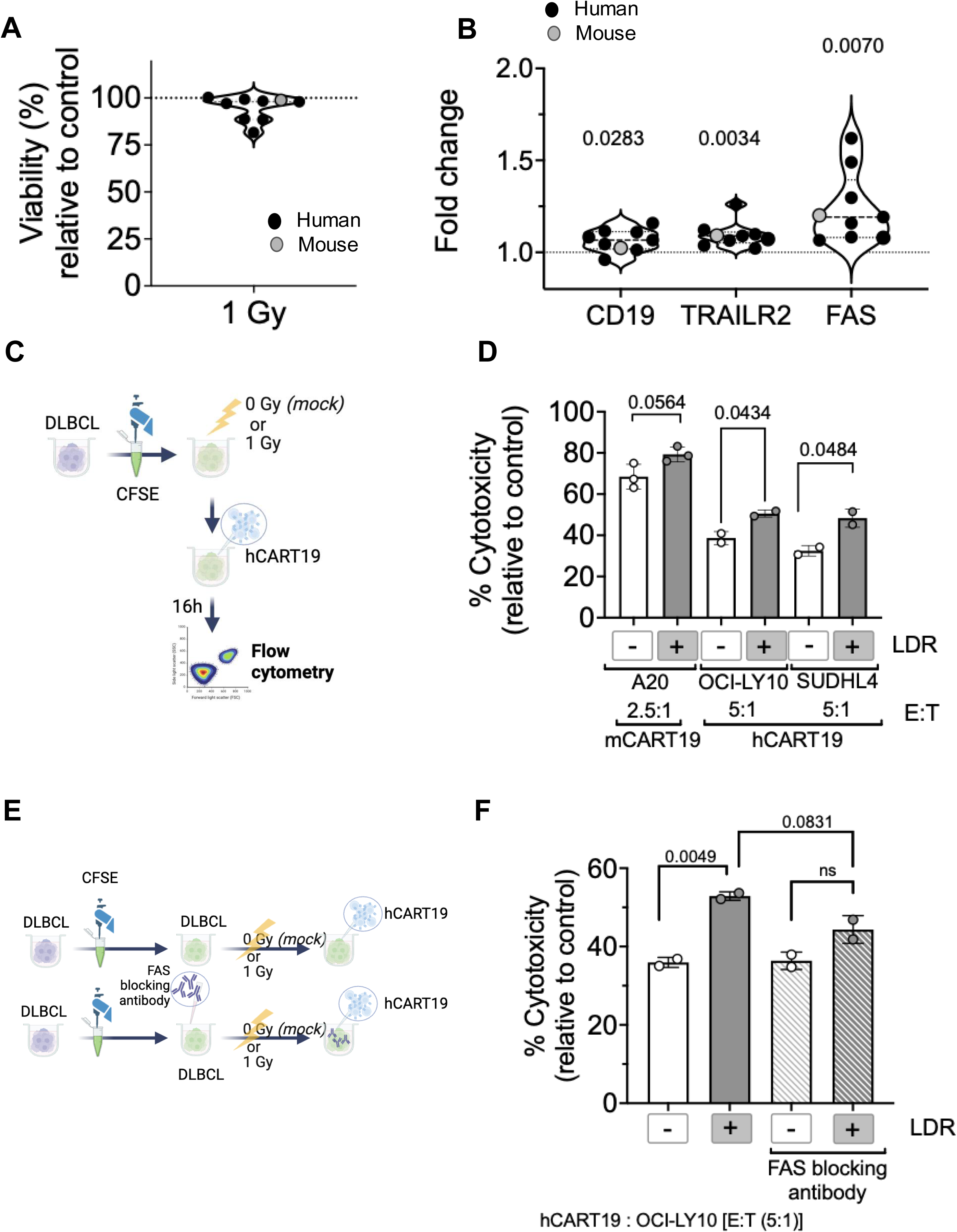
Radiation upregulates DRs in LBCL cells, enhancing the anti-lymphoma efficacy of CART19. **A.** Cell viability of eight human LBCL cell lines (OCI-LY10, SUDHL-2, RC-K8, SUDHL-4, SUDHL-5, SUDHL-6, OCI-LY1, and OCI-LY4), and a mouse-derived B cell lymphoma (A20 GFP-LUC); 24 hours post-LDR at 1 Gy, measured by flow cytometry. Each dot represents the average of at least 2 independent experiment per cell line performed at least in duplicate. **B.** Fold change of MFI for FAS, TRAIL-R2, and CD19 antigen (quantified by measuring antigen binding capacity (ABC) using BD Quantibrite^TM^ PE beads) 24 hours after irradiation with 1Gy. Each dot represents the average of at least 2 independent experiment per cell line performed at least in duplicate. Significance was calculated relative to mock irradiated control using paired t-test. **C.** Graphical representation for the experimental setup showing CFSE-labeled target cells (lymphoma cells) were irradiated at 1Gy or mock-irradiated, cells were then co-cultured with effector cells (hCART19) at defined E:T ratios. Cytotoxicity was assessed by flow cytometry using DAPI. **D.** Cytotoxicity of hCART19 against OCI-LY10 and SUDHL-4 and mCART19 against A20 GFP-LUC at the indicated E:T ratios, each dot represents the average of independent experiments performed in duplicate. Significance was calculated relative to mock irradiated control using an unpaired t-test. Plot depicts the mean and SD. **E.** Graphical representation for the experimental setup using a FAS blocking antibody. OCI-LY10 cells were stained with CFSE and incubated with FAS blocking antibody (2 μg/ml) for 1 hour before irradiation at 1Gy. Target cells were then cultured with effector cells (E:T= 5:1). The percentage of hCART19 cytotoxicity was calculated relative to the control (target cells at 0 (mock) or 1Gy). **F.** CART19 cytotoxicity assay performed with OCI-LY10 (with or without FAS blocking antibody), each dot represents the average of independent experiments performed in duplicate. Significance was calculated using unpaired t-test. Plot depicts the mean and SD.

As we observed the radiation-induced increase in CD19 antigen density and an increase expression in DRs, we hypothesized that such changes could improve CART19 cell anti-tumor efficacy. To this end, we manufactured human CART19 (hCART19) [17, 29], and mouse CART19 (mCART19) [30, 31], (**supplementary Fig. 2A, 2B, and 2C**), and assessed their cytotoxic capacity targeting CD19^+^ cells using human and mouse LBCL cell lines by performing *in vitro* co-culture assays (**Fig. 1C**). We found that hCART19 displayed significantly enhanced cytotoxicity when exposed to lymphoma cells irradiated with 1Gy dose compared to mock irradiated controls (**Fig. 1D, supplementary Fig. 1C)**. Similar to the effect on human lymphoma cells, 1Gy irradiated A20 GFP-LUC cells were more vulnerable to mCART19 compared to mock-irradiated controls **(Fig. 1D, supplementary Fig. 2D)**.

**Figure 2.**
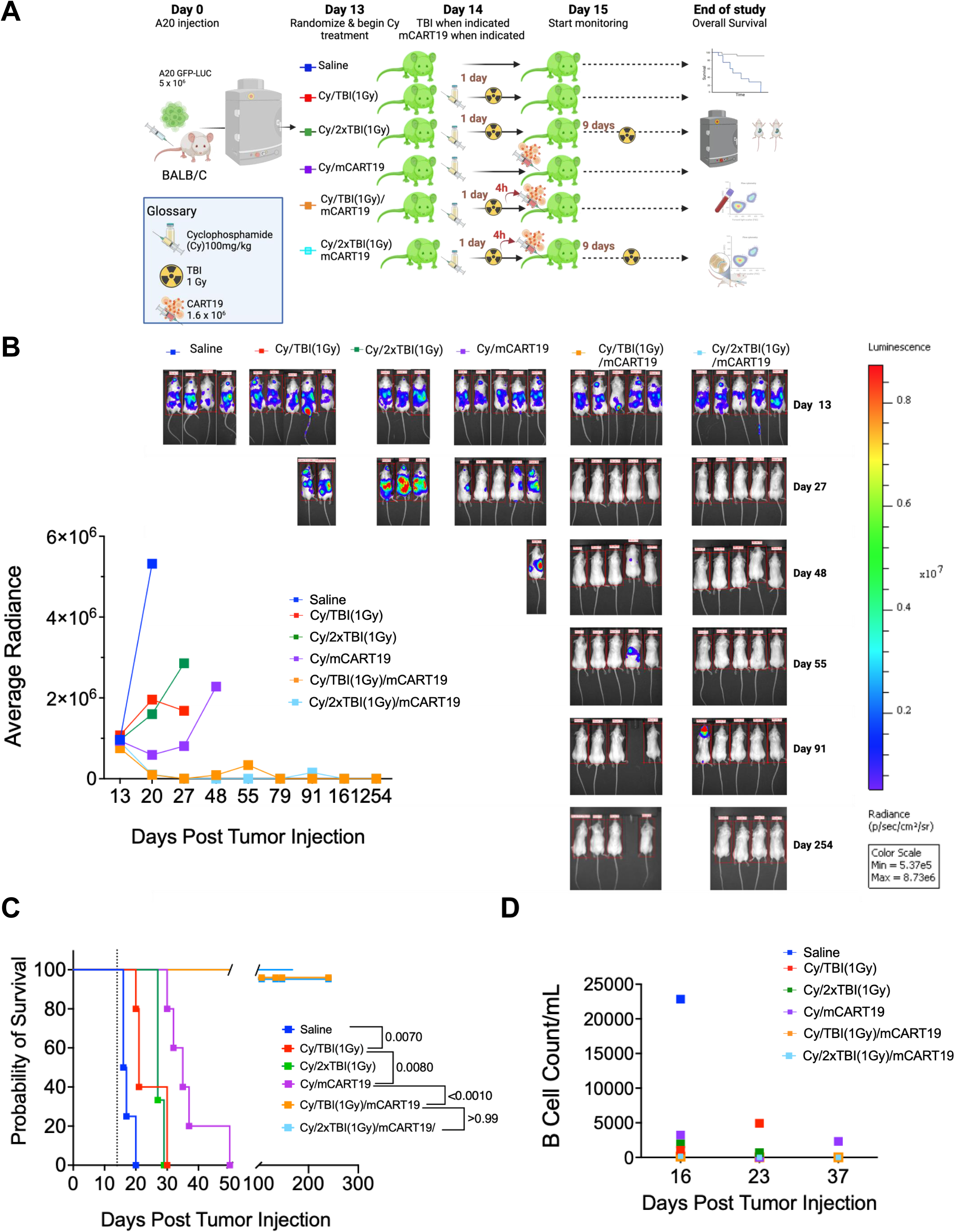
TBI (1Gy) enhances the efficacy of mCART19 in a LBCL syngeneic mouse model. **A.** Experimental layout for the syngeneic mouse model. BALB/c mice were transplanted with A20 GFP-LUC. Lymphoma (A20 cells) engraftment was confirmed by bioluminescence imaging (BLI) 13 days after injection. Animals were distributed into six cohorts as indicated 1: Saline control (n=4); 2: Cy/TBI(1Gy) (n=5); 3:Cy/TBI(1Gy) on day 14 and day 23 (n=3), 4:Cy/mCART19 (n=5), 5:Cy/TBI(1Gy)/mCART19 (n=5), and 6:Cy/TBI(1Gy) on day 14 and day 23, and mCART19 (n=5). TBI(1Gy) was given 4 hours before mCART19 injection to the indicated cohorts. Lymphoma (A20) progression was monitored weekly using BLI system. Bone marrow aspirates and peripheral blood draws were performed at multiple time points. **B. Left.** Average radiance (p/sec/cm2/sr) representing lymphoma growth at the indicated times (days). Each dot represents the mean per cohort. **Right.** BLI images for representative time points shown for all groups. **C.** Overall survival. Each square represents one animal*. P*-value calculated using Log-rank test**. D.** B cell count (absolute number of cells/mL) in peripheral blood at the indicated times (days). Each dot represents the mean of the surviving mice per group.

Since FAS was the most upregulated DR, we evaluated the potential role of FAS modulation on CART19 efficacy. To this end, we performed a cytotoxicity assay for human CART19 against OCI-LY10 cells (2.5:1 E:T ratio) in the presence or absence of a FAS-blocking antibody (**Fig. 1E**). OCI-LY10 cells were incubated with FAS blocking antibody before irradiation and then co-cultured with CART19. CART19 cytotoxicity was significantly reduced (albeit not abrogated) when the FAS-blocking antibody was added to irradiated OCI-Ly10, indicating a role for FAS (**Fig. 1F**).

Collectively, our results underscore the capacity of 1Gy radiation to increase CD19 antigen density and the expression of DRs in both human and mouse LBCL cells *in vitro* and significantly enhance the cytotoxic capacity of CART19 against lymphoma.

### TBI at 1Gy augments the therapeutic efficacy of mCART19 in an immunocompetent syngeneic LBCL mouse model

To determine whether the same 1Gy dose of radiation can also improve the activity of CAR T cells *in vivo*, we employed a syngeneic mouse model, this model enables the implementation of a lymphodepletion regimen similar to what patients receive in clinical setting [30]. Briefly, BALB/c mice were intravenously injected with 5X10^6^ A20 GFP-LUC cells; 13 days later, tumor burden was evaluated using bioluminescence imaging (BLI) to corroborate lymphoma engraftment and assign animals to treatment cohorts. The lymphodepletion regimen consisted of 100 mg/kg cyclophosphamide (Cy), a dose that resulted in effective lymphodepletion and was confirmed not to cure the mice (**supplementary Fig. 3A**), consistent with a prior study [30]. In addition to evaluating the benefit of single 1Gy dose of TBI before mCART19, we also tested delivering a second dose of TBI (1Gy) nine days after mCART19 treatment in an additional experimental cohort of mice. Animals were assigned to the following cohorts on day 13: 1) Saline; n=4, 2) Cy/TBI(1Gy) on day 14 post tumor-injection; n=3; 3) Cy/mCART19 n=5; 4) Cy/TBI(1Gy) on day 14/mCART19 n=5, 5) Cy/2xTBI(1Gy), TBI was given on day 14 and day 23, n=5; and 6) Cy/2xTBI(1Gy)/mCART19, TBI was given on day 14 and day 23, n=5 (**Fig. 2A**). Cy (100 mg/kg) was administered intraperitoneally (i.p) 24 hours before mCART19 treatment. TBI at a dose of 1Gy was delivered as a single fraction 4 hours before mCART19 administration, or 9 days later after mCART19 for the animals that received 2 doses. mCART19 cells were delivered intravenously at a dose of 1.6X10^6^ CART cells. As expected, Cy/mCART19 improved overall survival (OS) compared to Cy/TBI(1Gy) (median OS= 35 days and 21 days, respectively; *p*=0.0080). Strikingly, 4/5 of the mice treated with Cy/TBI(1Gy)/mCART19 and 4/5 of the mice receiving Cy/2xTBI(1Gy)/mCART19/ remained alive and disease-free at eight months post-treatment (*p*<0.001), with median OS not reached **(Fig. 2B, 2C)**. In the Cy/TBI(1Gy)/mCART19 group, one mouse succumbed to a florid relapse on day 59, while in the Cy/2xTBI(1Gy)/mCART19 group, one mouse died from an isolated CNS relapse on day 91. Of note, all relapses were CD19-positive. As an indirect biomarker of mCART19 activity, absolute B cell counts in the peripheral blood (PB) were monitored. We found that mice that received mCART19 exhibited B cell aplasia (**Fig. 2D, supplementary Fig. 3B**). Since one dose of 1Gy TBI before mCART19 in this initial experiment was very effective, the potential benefit of the second dose of 1Gy TBI was not evaluable. Our results revealed that *in vivo* radiation exerts a more pronounced effect than *in vitro*, indicating additional mechanisms by which 1Gy dose of TBI augments the efficacy of mCART19.

### TBI at a dose of 1Gy increases *in vivo* expression of FAS, augments lymphodepletion, enhances mCART19 expansion and increases CCL2 chemokine

We next sought to identify the mechanisms underlying this radiation effect in *vivo*. Since our *in vitro* studies showed that FAS was the most upregulated DR, we first measured the impact of TBI (1Gy) on the expression of FAS in A20 lymphoma cells *in vivo*. We observed a 1.5-fold increase in FAS expression on lymphoma cells isolated from the liver of irradiated mice when compared to the mock-irradiated/saline cohort (*p*=0.0169) (**Fig. 3A**). These data suggest that FAS is an important component of the priming achieved by TBI (1Gy). As the responses we observed were more striking *in vivo* than *in vitro*, we investigated additional effects of TBI(1Gy). Studies have shown that intensified depletion of endogenous lymphocytes before adoptive transfer of CART19 enhances its peak of expansion and persistence, leading to improved CR rate, progression-free survival, and OS [32, 33]. Thus, we tested whether using TBI at a dose of 1Gy after the conditioning regimen (Cy) would affect the depth of lymphodepletion. To this end, we placed immunocompetent mice (BALB/c) into four cohorts: saline, Cy (100 mg/kg), TBI (1Gy) only, and a combination of Cy and TBI(1Gy) (**supplementary Fig. 4A**). Mice that received the combination of TBI(1Gy) and Cy had a significantly decreased absolute lymphocyte count three days post-treatment compared to those who received Cy only (*p*<0.001) and/or TBI(1Gy) only suggesting enhanced lymphodepletion with the combination (**Fig. 3B**). Importantly, all groups recovered to a normal baseline lymphocyte count by day 60, with a kinetic of recovery of lymphopenia comparable to healthy controls exposed to Cy or 1Gy TBI. Notably, while the extent of lymphodepletion deepened, there was no significant difference in neutropenia between the groups that received Cy or the combination of Cy and TBI (1Gy), and all groups recovered neutrophil count by day 60 (**Fig. 3B**).

**Figure 3.**
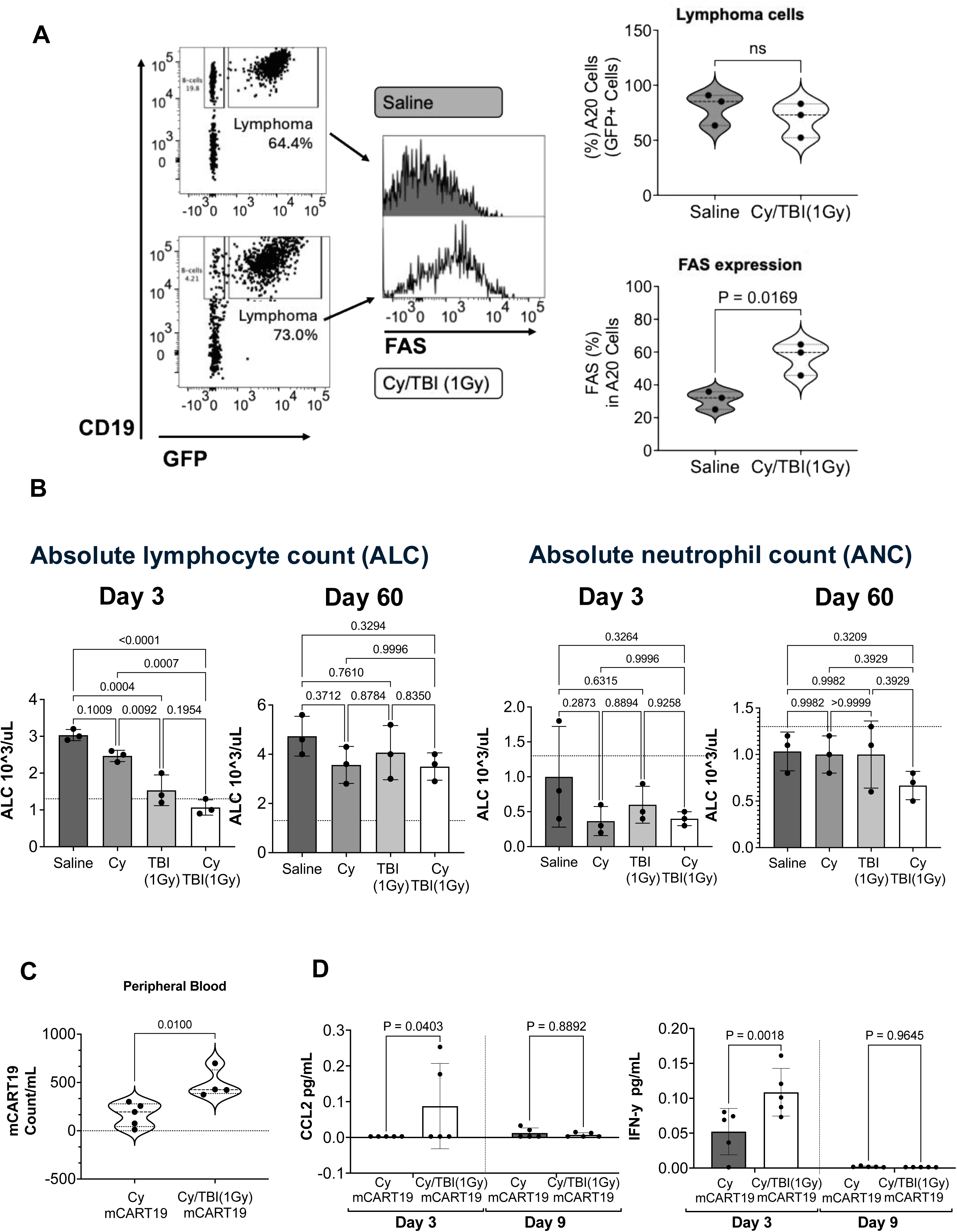
TBI (1Gy) upregulates FAS *in vivo*, enhances lymphodepletion and mCART19 expansion, increasing the cytokine release. A. Left. Representative dot plots for the gating of A20 GFP-LUC lymphoma cells (GFP^+^) isolated at day 3 post-treatment initiation from the liver of mice that received Cy/TBI(1Gy) vs. saline control. **Right (top).** Percentage of lymphoma (A20) cells in mice that received Cy/TBI(1Gy) vs. saline control. **(bottom)** Percentage of FAS expression in lymphoma cells in the indicated groups: Cy/TBI(1Gy) vs. saline control, 3 days post radiation. Each dot represents one animal*. P*-value calculated using unpaired t-test. Violin plots depict the median and quartiles. **B.** Lymphocyte and neutrophil count on day 3 and day 60 post-treatment in healthy mice exposed to the indicated treatments. The dotted line represents the lower limit of the normal range of blood cells: 1.3 10^3^/µL for lymphocytes and 0.4 10^3^/µL for neutrophils. Each dot represents one animal, the bar represents the mean with SD. *P*-value calculated using one-way ANOVA. **C.** mCART19 (count/mL) from peripheral blood of mice 9 days post-treatment initiation comparing the groups that received mCART19 with or without TBI(1Gy). Each dot represents one animal*. P*-value calculated using unpaired t-test. **D.** CCL2 and IFN-γ concentrations in the plasma of mice on day 3 and day 9 post-mCART19 treatment initiation for the conditions receiving mCART19. Each dot represents one animal, and the bar represents the mean with SD.

In addition to changes in lymphodepletion, we investigated the changes in CAR T cell counts. Several studies indicated that post-infusion CART19 cell counts after infusion are critical in predicting response durability, and the detection of an early peak expansion was associated with improved clinical responses in LBCL [1, 2, 34–36]. Thus, we used the syngeneic LBCL mouse model to evaluate the effect of TBI(1Gy) on mCART19 expansion (**supplementary Fig. 4B**). We observed a 2.8-fold increase of mCART19 on day 9 in the PB of mice receiving Cy/TBI(1Gy)/mCART19 when compared to the Cy/mCART19 cohort (*p*=0.0100) (**Fig. 3C, supplementary Fig. 4C**).

Since we detected an increase in mCART19 expansion, we next sought to assess changes in cytokines that may mediate this effect. We evaluated the 15 cytokines using plasma from PB at day 3 and day 9, using a panel that included: IFN-γ, IL-10, CCL4 (MIP-1β), IFN-α, CXCL9 (MIG), CXCL10 (IP-10), TNF-α, IL-6, VEGF, IL-4, IL-18, IL-1b, CCL3 (MIP-1α), CCL2 (MCP-1), and GM-CSF. On day 3 post-treatment, we found a significant increase in CCL2 (also known as monocyte chemoattractant protein-1 (MCP-1)) in the Cy/TBI(1Gy)/mCART19 cohort (*p*=0.0403) (**Fig. 3D**). CCL2 is a chemokine that plays a critical role in attracting CAR T cells to tumor sites, enhancing their ability to infiltrate the tumor microenvironment and effectively destroy tumor cells [37–43]. Consistent with anti-tumor activity in the Cy/TBI(1Gy)/mCART19 group, we found IFN-γ also significantly elevated (*p*=0.0018) (**Fig. 3D**). CXCL10 and CXCL9 were higher on day 3 in Cy/TBI(1Gy)/mCART19, but the increase did not reach statistical significance (**supplementary Fig. 4D**). This generalized cytokine surge was accompanied by clinical features associated with cytokine release syndrome (CRS). A clinical body score has been developed to assess mCART19 effect in mice [44]. We adopted a similar score to evaluate mice for hunched posture, ruffled fur, and weight loss (1 point each, **supplementary table 1**). Mice that received mCART19 began to exhibit CRS-like symptoms by day two post-mCART19 administration, which resolved by day 7. Mice that also received TBI (1Gy) scored higher for CRS-like features (*p*=0.0036) (**supplementary Fig. 4E)**; however, importantly, this event did not translate into fatal toxicities.

Taken together, our results in a syngeneic mouse model confirmed the ability of radiation to upregulate FAS on tumor cells *in vivo*, as we had observed *in vitro* in the setting of enhanced CART19 cytotoxicity. Furthermore, the data demonstrate that TBI (1Gy) deepens lymphodepletion, promotes substantial mCART19 expansion in peripheral blood, and increases the secretion of IFN-γ and CCL2 without significant toxicity or prolonged cytopenia.

### TBI at a dose of 1Gy boosts mCART19 intra-tumoral trafficking capacity and improves persistence

Since CCL2 has been reported to be a key driver for CAR T cell trafficking to tumors and given our observation that CCL2 significantly increased after TBI (1Gy), we hypothesized that TBI results in increased trafficking of CAR T cells to lymphoma infiltrating sites. Thus, we established four cohorts of mice bearing LBCL (A20 GFP-LUC) to receive the following treatments: Cy/TBI(1Gy), Cy/mCART19, Cy/TBI(1Gy)/mCART19 and saline only (**Fig. 4A**). As in all other experiments, 1Gy TBI was given 4 hours before mCART19 treatment (1.6x10^6^ cells/mouse). Three days after mCART19 administration, the spleen, liver, and bone marrow were harvested and processed for flow cytometry analysis. We found that TBI (1Gy) resulted in a of 2.7-fold and 1.7-fold increase in mCART19 cell counts within the lymphoma infiltrating organs such as the spleen and the liver respectively [30], relative to the animals treated just with Cy prior mCART19 treatment (*p*=0.0284 and *p*=0.0169, respectively), suggesting that TBI (1Gy) markedly improved the tumor-trafficking capability of mCART19 (**Fig. 4B**). Although there was 1.6-fold increase in mCART19 in the bone marrow in the group that received Cy/TBI(1Gy)/mCART19, it did not reach significance (*p*=0.2758) when compared to Cy/mCART19 group as observed for the lymphoma infiltrating organs (**supplementary Fig. 5A**).

**Figure 4.**
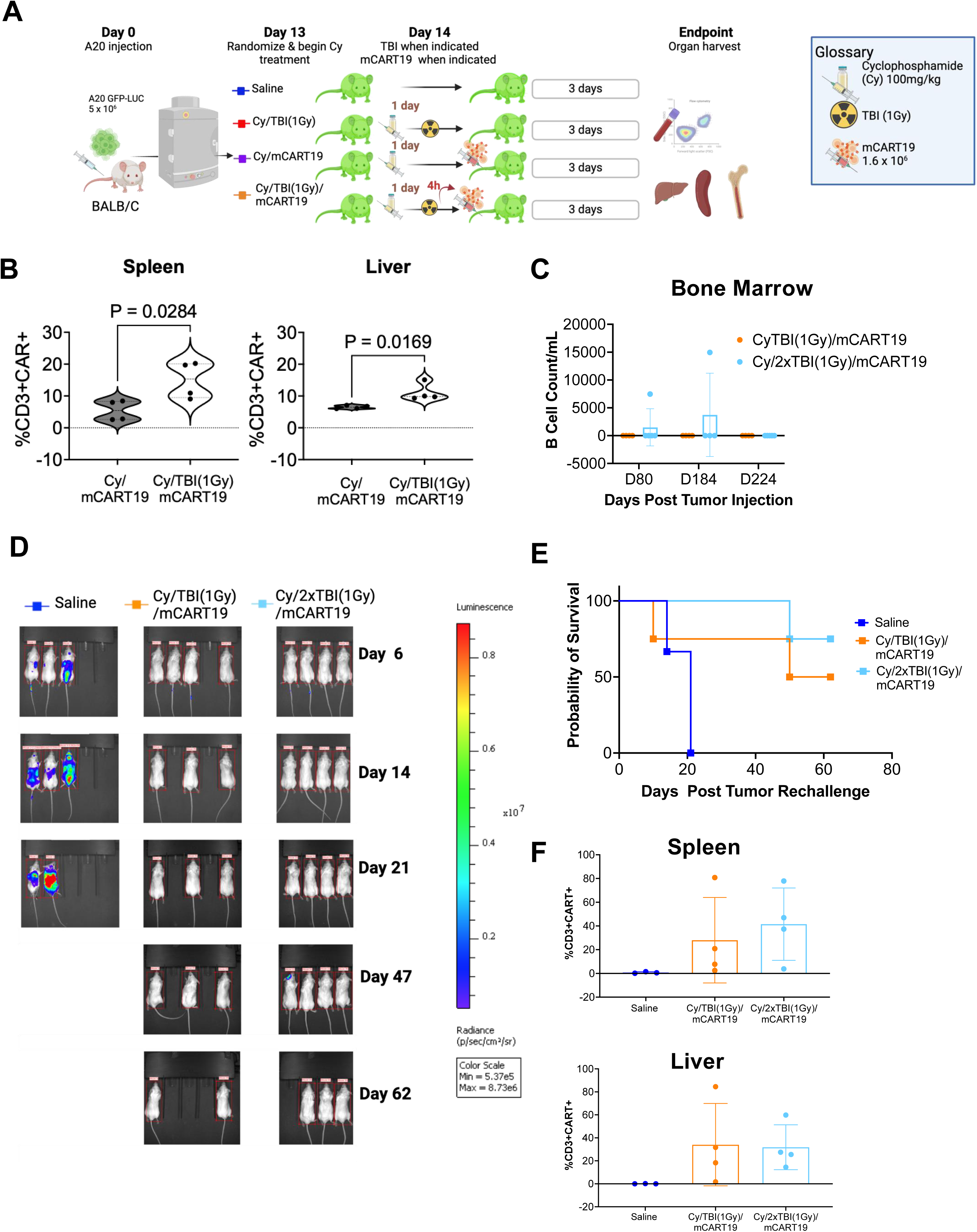
TBI (1Gy) improves CART19 intra-tumoral trafficking and persistence. **A.** Graphical representation of the experimental setup for the evaluation of mCART19 trafficking. BALB/c mice were I.V. injected with A20 GFP-LUC. Lymphoma (A20) engraftment was confirmed by BLI 13 days after injection. Cohorts were assigned as indicated: 1: Saline (n=3), 2: Cy/TBI(1Gy) (n=3), 3: Cy/mCART19 (n=4), 4: Cy/TBI(1Gy)/mCART19 (n=4). TBI(1Gy) was given 4 hours before mCART19 injection. Organs were harvested on day 3 post-mCART19 treatment initiation and cells were evaluated by flow cytometry. **B.** mCART19 (CD3^+^ and Myc-tag^+^ T cells) percentage in liver and spleen at day 3 post-mCART19 treatment (gating strategy shown in **supplementary Fig.4**). Violin plots depict the median and quartiles, each dot represents one animal*. P*-value calculated using unpaired t-test. **C.** B cell aplasia in long-term surviving mice that received TBI(1Gy) and mCART19 from Fig 2. Graph shows the number of B cells (CD19+GFP-) (count/mL) in the BM (obtained by BM aspirates at the indicated days post lymphoma injection. Each dot represents one animal surviving, and the bar represents the mean with SD. **D.** Bioluminescence images for the surviving mice from Fig 2 treated with Cy/TBI(1Gy)/mCART19 (n=4) and Cy/2xTBI(1Gy)/mCART19 (n=4) that were rechallenged with A20 GFP-LUC at day 274 post initial tumor injection. A new group of BALB/c mice (n=3) were injected with A20 GFP-LUC as controls. Representative images are shown from weekly assessment of BLI. **E.** Overall survival post rechallenge. Each square represents one animal. **F.** Animals were sacrificed at 62 days post rechallenge, percent of mCART19 in harvested organs (spleen and liver) is shown, each dot represents one mouse, and the bar represents the mean with SD.

We hypothesized that improved persistence of CAR^+^ cells was another mechanism of improved OS in mice receiving combined TBI(1Gy) and mCART19. Thus, to confirm and assess the persistence of mCART19 cells, we evaluated B cell aplasia as a surrogate marker in the long-term surviving mice from the 1Gy dose TBI exposure (4 mice in Cy/TBI(1Gy)/mCART19 and 4 mice in Cy/2xTBI(1Gy)/mCART19) by performing bone marrow aspirates at days 80, 184, and 224 post-tumor cell injection. We found that the surviving mice maintained complete B cell aplasia, confirming sustained functional mCART19 persistence for more than six months after administration (**Fig. 4C**). One animal appeared to have B cell recovery at day 80, despite remaining in remission. However, by day 274, that animal had no measurable B cells and remained disease-free. To further confirm functional mCART19 persistence and its ability to protect against relapse, we rechallenged the surviving mice from the Cy/TBI(1Gy)/mCART19 and Cy/2xTBI(1Gy)/mCART19 cohorts, previously described in **Fig. 2**, with A20 GFP-LUC cells, at day 274 post initial tumor injection. To ensure the quality of the engraftment, we injected A20 GFP-LUC cells into a new cohort of BALB/c mice as a control for this experiment. Seven of eight surviving mice that received 1Gy dose of TBI and mCART19 rejected tumor rechallenge, demonstrating sustained protection conferred by mCART19. In contrast, control mice succumbed to disease as expected (**Fig. 4D, 4E, supplementary Fig. 5B**).

One mouse from Cy/2xTBI(1Gy)/mCART19 died of isolated CNS relapse after tumor rechallenge. Two mice from Cy/TBI(1Gy)/mCART19 had to be euthanized, however, these animals did not appear to die from disease relapse as there was no evidence of A20 GFP-LUC cells on imaging or upon processing their organs by flow cytometry. These deaths likely reflected natural aging effects in animals that were 15-16 months old [45]. All surviving mice were euthanized 62 days post-rechallenge. Spleen and liver were harvested to assess the presence of mCART19 cells and to evaluate for any potential residual disease. mCART19 cells were clearly present in all these organs 336 days post-mCART19 injection (**Fig. 4F**).

Our results provide evidence that TBI at a dose of 1Gy exerts multi-faceted effects on mCART19, including promoting persistence and enhancing intra-tumoral migration.

### TBI at a dose of 1Gy impacts T cell and mCART19 subsets, with differentially expressed genes related to early activation, cytotoxicity, and proliferation

To further elucidate the impact of TBI(1Gy) on T cell and mCART19 repertoires and activity against LBCL, we conducted single-cell RNA sequencing (scRNA-seq) to compare the profiles of lymphoma-bearing mice treated with Cy/mCART19 to those with Cy/TBI(1Gy)/mCART19. Profiles were evaluated 3 days post-treatment (**Fig. 4A**). First, we clustered all cell types using commonly deployed analytical methods and visualized resultant clusters using uniform manifold approximation and projection (UMAP) (**Fig. 5A, supplementary Fig. 6B**). The mice in Cy/TBI(1Gy)/mCART19 group exhibited an increase in total T cells accompanied by a significant decrease in B cells compared to the Cy/mCART19 (**Figure 5A and 5B**), confirming that TBI at a dose of 1Gy improves T cell expansion and elimination of CD19^+^ cells. Importantly, these data show that TBI(1Gy) also affects autologous T cells that do not have the CAR transgene but may have anti-tumor capacity. Further clustering T cell subsets, we identified eleven distinct phenotypes, ten of them CD4^+^ and one CD8^+^ (**Fig. 5C, supplementary Fig. 6C**). We identified a preferential expansion of almost all T cell subsets in Cy/TBI(1Gy)/mCART19 mice compared to Cy/mCART19 group, with the most notable increase in CD4^+^ effector cytotoxic (eCTL) subset (**Fig. 5C**), a population found to play a key role in tumor regression of mouse lymphoma [46]. This subset was recently recognized for its potent anti-tumor properties [47, 48]. Conversely, the regulatory T cells subsets remained unchanged (**Fig. 5C**). Studies have suggested that less-differentiated naive and memory CD4^+^ T cells tend to persist longer and provide more potent anti-tumor activity [49, 50] in both lymphoma [51] and leukemia [52]. Next, we performed differential expression analysis (DEA) of genes of T cell subsets to determine whether there was a potential difference within the same subsets under different conditions. A previous report emphasized the crucial role of CD8^+^ T cells in the immune response to radiotherapy [6, 8]; we found that CD8^+^ T cells in mice that received 1Gy dose of TBI upregulated genes and pathways related to IFN-γ signaling, T cell proliferation, and immune-modulatory properties in the tumor microenvironment (**Fig. 5D, 5E**), suggesting enhanced CD8^+^-mediated pro-inflammatory and anti-tumor characteristics [53–55]. In addition, DEA and Gene set enrichment analysis showed that the Naive_1 subset in Cy/TBI(1Gy)/mCART19 upregulated genes and pathways linked to T cell proliferation, mitochondrial oxidative phosphorylation, protein production, and IFN-γ response (**supplementary Fig. 6A**), [53, 55].

**Figure 5.**
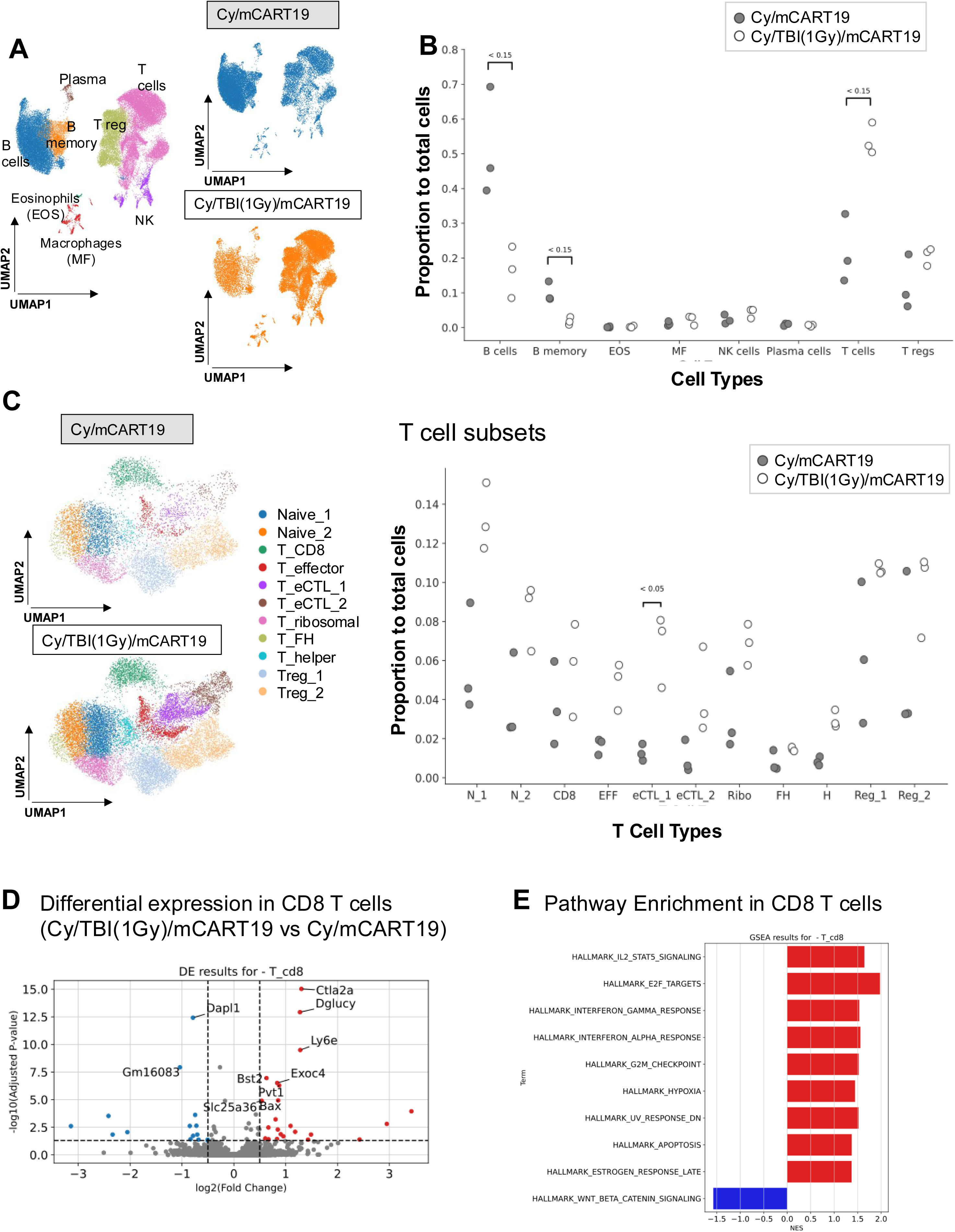
TBI (1Gy) influences T cell differentiation, enhancing subsets with superior anti-tumor characteristics. A. Left. UMAP of scRNA-seq data of cells in the dataset from spleens from mice at the indicated conditions (Cy/mCART19 and Cy/TBI(1Gy)/mCART19 day 3 post-mCART19 treatment initiation) annotated by cluster (color). **Center.** UMAP embeddings showing the distribution of immune cells in clusters by condition, Cy/mCART19 (24,556 cells) and Cy/TBI(1Gy)/mCART19 (24,412 cells). **B.** Dot plot showing the proportion of immune cell types to a total dataset of cells. Each plot represents the cell type within a condition, and each circle represents the proportion per mouse. (EOS: eosinophils; MF: macrophages; NK: natural killer). Changes in the proportion of cells between groups were calculated based on compositional data analysis with an adjusted *P*-value is shown**. C. Left.** UMAP of scRNA-seq data of T cell subsets in the dataset sorted from spleens of BALB/c mice in both conditions (Cy/mCART19 and Cy/TBI(1Gy)/mCART19) annotated by cluster (color). **Right.** Dot plot showing the proportion of T cells within each subcategory of T cells to the total dataset of cells. Each plot represents the subset of T cells within the condition, and each circle represents a mouse. Changes in the proportion of cells between groups were calculated based using compositional data analysis. **D.** Volcano plot of Wilcoxon rank-sum scores from the differential gene expression analysis of Cy/TBI(1Gy)/mCART19 vs Cy/mCART19. in CD8^+^ T cells. Genes with adjusted *P*-value=1 were filtered out before plotting. **E.** Top 10 enriched pathways of CD8^+^ T cell subset in Cy/TBI(1Gy)/mCART19 vs. Cy/mCART19.

Interestingly, when analyzing subsets of the CAR^+^ T cells, we found a significant increase in CD8^+^ T cells and a reduction in T regulatory cells in the group that received 1Gy dose of TBI (**Fig. 6A**). Previous studies have indicated that CAR^+^ Tregs are implicated in hindering CART19 efficacy and associated with disease progression [56]. These data suggest that mCART19 cells in the irradiated group may be the best fit to eliminate tumor cells with less hindrance from CAR^+^ Tregs. It is important to note that there was no significant difference in Treg clusters when comparing all T cells in both groups (**Fig. 5C).** Thus, the changes we observed are preferential to CAR^+^ Tregs. Consistently, mCART19 in mice that received 1Gy dose of TBI showed a striking downregulation of genes associated with T cell inhibition and regulation while simultaneously upregulating multiple genes and pathways critical for early T cell activation, cytotoxicity, proliferation, and cell adhesion, when compared to the Cy/mCART19 group (**Fig. 6B**) [57].

**Figure 6.**
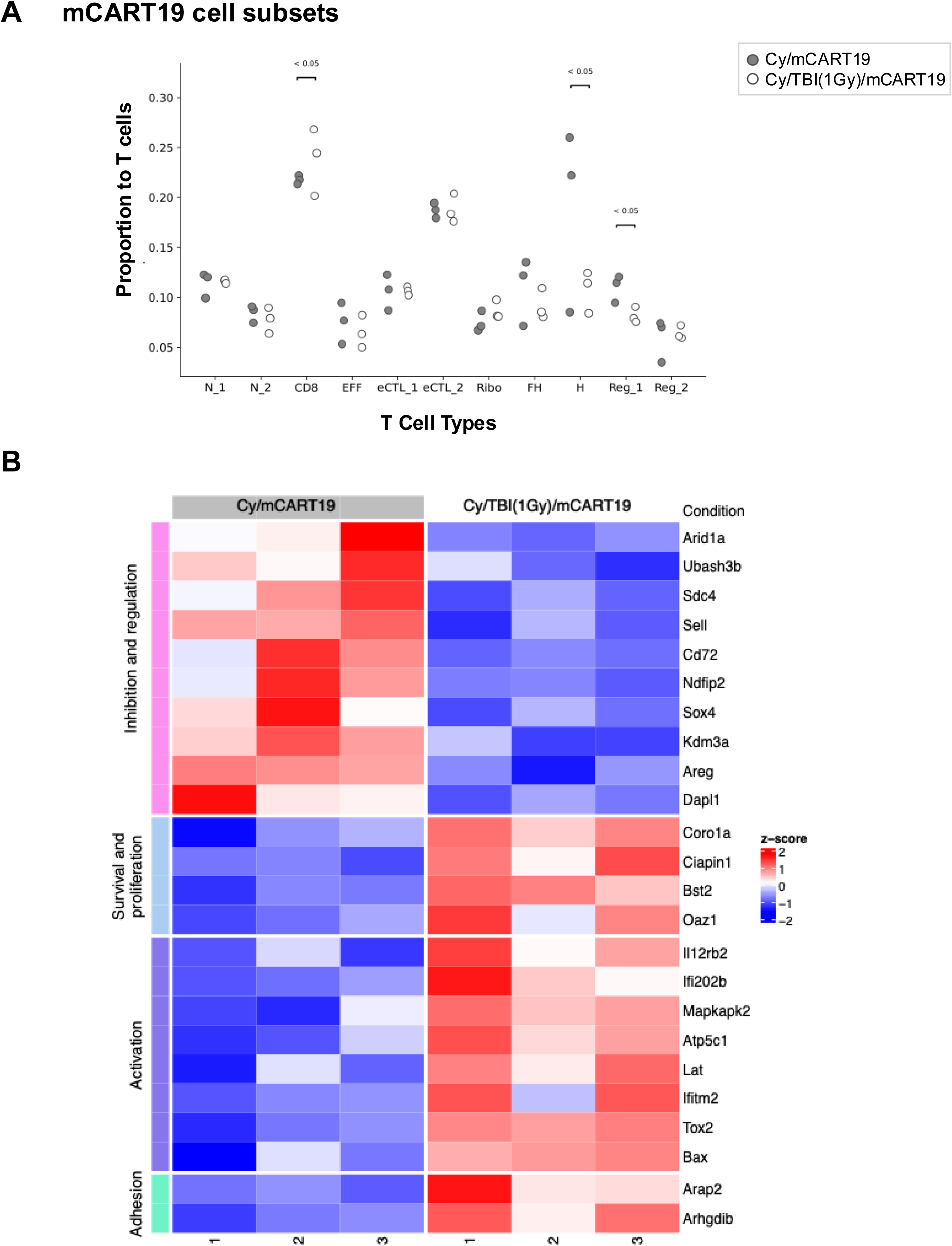
LD-TBI influenced mCART19 differentiation, leading to the upregulation of genes involved in early activation, cytotoxicity, and proliferation. **A.** Dot plot showing the proportion of CAR^+^ T cells within each subcategory of CAR^+^ T cells. The Subset of CAR^+^ T cells is shown within the indicated condition, and each circle represents a mouse. Changes in the proportion of CAR^+^ T cells between groups were calculated using a cell type-specific binomial test. **B.** Heatmap of expression of selected markers found dysregulated under padj < 0.05 in mCART19 cells between conditions. Each cell in the heatmap represents the z-scored transformed expression of the gene in one mouse per condition.

Taken together, the transcriptional studies support the *in vitro* and *in vivo* observations of a favorable impact of TBI (1Gy) to improve the anti-tumor activity of both T cells and mCART19 adoptively transferred to the lymphoma-bearing mice.

## DISCUSSION

Despite the notable success of CART19 therapy in lymphoma patients, insufficient or transient responses remain significant challenges. These hurdles have been attributed to several factors, broadly grouped into three categories: i) tumor-intrinsic factors, related to underlying disease biology and genomic composition such as impaired death signaling pathway or low antigen density [28, 58, 59]; ii) immunosuppressive tumor microenvironment, which impedes intra-tumoral trafficking and functionality of CART19 [28, 60], and iii) CAR product-related factors, including suboptimal expansion kinetics [1, 2, 34, 36, 61] and an unfavorable phenotypic profile, with a higher proportion of CAR^+^ Treg cells and fewer CAR^+^ naive and memory T cells [51, 56, 62–64]. Addressing these resistance mechanisms remains a critical step to advancing and improving the therapeutic efficacy of CART19. Here, we demonstrate that the use of TBI at 1Gy can overcome many of the above-mentioned hurdles.

Using both *in vitro* and *in vivo* approaches, we showed that 1 Gy radiation improves CART19 anti-lymphoma activity *in vitro*, and we implemented such a strategy *in vivo,* in the form of TBI (1Gy), noting a striking therapeutic effect against LBCL. Our studies revealed that TBI delivered as a single 1Gy dose, a highly tolerable regimen, exerts the multifaceted effects needed to prevent and potentially overcome obstacles to CAR T cell therapy success. Specifically, in our model, 1Gy dose of TBI improves lymphodepletion, augments the expression of DRs and CD19 antigen density in lymphoma cells, affects subsets distributions of T cells and CAR^+^ T cells, and allows improved trafficking to tumor sites. Our findings expand on prior work by our group [17] and Kim et al. [65] that showed a role of ionizing radiation in augmenting the efficacy of CART19 cells targeting ALL in immunodeficient mouse models. While we cannot exclude the potential immunogenic effect of GFP, that could result in protection against the lymphoma cells (A20 GFP-LUC) during the rechallenge experiment, we can conclude that to have a protective effect the mCART19 have to be present. Persistence of the CAR T cells was confirmed after the rechallenge experiment reinforcing this hypothesis. Because of the important clinical implications of our observations, we will pursue future experiments to further validate the long term protection that TBI(1Gy) has provided beyond CAR T persistence. For example, rechallenging mice with A20-LUC cells that do not express GFP, and with A20-LUC cells lacking CD19 expression [66].

Beyond the demonstrated benefits of 1Gy dose of TBI on CART19 expansion, persistence, early activation, and phenotypic composition, the enhanced lymphodepletion achieved by combining 1Gy dose of TBI with the standard regimen further supports its integration into the conditioning protocol for CART19 therapy. Unlike other forms of localized radiation, TBI (1Gy) provides systemic exposure to lymphoma cells, which is particularly advantageous for addressing the disseminated nature of the disease. In addition, a single dose of 1Gy TBI well tolerated and historically was used for treatment of refractory lymphoma [67–69]. This is especially relevant for high-risk patients with high tumor burden or measurable disease at the time of CART19 therapy, where outcomes are generally less favorable [70, 71]. TBI at a dose of 1Gy could also be tested in other conditioning protocols for CAR T cell therapies targeting various solid tumor models, where challenges such as CAR T cell trafficking into tumor tissues and long-term persistence are also observed [72].

We leveraged the compatibility of an immunocompetent syngeneic mouse model, which closely mirrors immune interactions in human, to assess the safety and preliminary efficacy of incorporating TBI at a dose of 1Gy with CART19 therapy. Prolonged cytopenia is of particular concern as radiation can impair bone marrow recovery [73]. Furthermore, immune effector cell-associated hematotoxicity (ICAHT) has been recognized as a significant complication of CAR T cell therapy [74, 75] and can predispose to infectious complications [76], which are the primary driver of non-relapse mortality in CAR T recipients [77, 78]. Thus, it was essential to demonstrate that a single TBI session at 1 Gy dose did not significantly impact hematologic recovery in our model. TBI (1Gy) did not result in fatal CRS despite the notable early activation and surge in pro-inflammatory cytokines. These findings are encouraging and have prompted a Phase I/II trial testing this paradigm in human.

Our results indicate that TBI (1Gy) rapidly impacts T cell subset differentiation, with changes observable as early as three days post-radiation. scRNA-seq data revealed an overall enrichment of T cells, particularly the CD4^+^ effector cytotoxic T cells. While less differentiated T cells possess anti-tumor efficacy [51, 52], CD4^+^ effector cytotoxic cells can produce IFN-γ and express granzymes and perforin, leading to a potent CD8^+^ T cell-independent tumor control and resulting in durable anti-tumoral memory responses [47, 48]. Our study demonstrates that the TBI (1Gy) effect extends beyond CAR^+^ T cells, and it is not limited to the CAR-target interaction. While CAR T cells are activated through continuous antigen exposure and co-stimulatory signals, we showed that 1Gy dose TBI amplifies this activation, which leads to a distinct molecular profile enriched in genes related to T cell activation and proliferation coupled with suppression of regulatory markers. The long-term effects of TBI (1Gy) on CART19 and the identification of specific CAR^+^ and T cell subsets contributing to sustained longevity and tumor control remain to be explored in future studies.

In conclusion, we demonstrate that utilizing TBI at a dose of 1Gy significantly enhances CART19 efficacy by improving CAR T cell expansion, persistence, intra-tumoral trafficking, and phenotypic composition. This approach provides a valuable foundation for further research and holds promise for integration into clinical practice to further improve CART19 outcomes in lymphoma patients.

## METHODS

### Cell lines

ABC-LBCL (OCI-LY10, SUDHL-2, and RC-K8) and GCB-LBCL (SUDHL-4, SUDHL-5, SUDHL-6, OCI-LY1, and OCI-LY4), were provided by Dr. Mark Minden (Princess Margaret, Toronto, Canada) and Dr. Ari Melnick (Weill Cornell Medicine, NY, USA) under a material transfer agreement (MTA). The NALM6 (human acute lymphoblastic leukemia) cell line was obtained from the American Tissue Culture Collection (ATCC, Manassas, VA, USA). A20 cell line (BALB/c mouse-derived B-cell lymphoma) transduced with a GFP-luciferase fusion gene (A20 GFP-LUC) was provided by Dr. Renier Brentjens (Roswell Park, NY, USA). All cell lines underwent authentication using short tandem repeat (STR) analysis (University of Arizona Genetics Core or Bio-Synthesis Inc.) and are routinely tested for *Mycoplasma spp.* with PlasmoTest^TM^ Mycoplasma Detection Kit (InvivoGen, #rep-pt1).

### Generation of human CART19 cells

Second generation CD19-targeting CAR construct consists of an extracellular antigen-binding domain (CD19-scFV), CD8 for hinge and transmembrane domain, 4-1BB co-stimulatory domain, and CD3ζ chain signaling domain followed by EGFRt as a tag. PBMC were collected from buffy coat from healthy donors. T cells were purified on LS columns (Miltenyi Biotec, # 130-042-401) using CD4 and CD8 microbeads (Miltenyi Biotec, #130-045-101 and #130-045-201), were activated with CD3/CD28 beads (T Cell TransAct; Miltenyi Biotec, #130-111-160) and incubated for 24 h. Then, activated T cells were transduced with lentiviral vector (pCDH-EF1a-CD19 (FMC63)-2nd(4-1BB)- EGFRt; Creative biolabs) carrying the h194-1BBz construct. Activated T cells were cultured in TexMACS media (Miltenyi Biotec, #130-097-196) with hIL-7 (155U/mL, Miltenyi Biotec, # 130-095-362) and hIL-15 (290U/mL, Miltenyi Biotec, #130-095-762). CAR positivity was confirmed by the expression of EGFRt in CD45^+^CD3^+^CD19^−^ (**Supplementary Table 3**) population by flow cytometry. Expanded CART19 cells were frozen and stored in vials in liquid nitrogen before use.

### Generation of retroviral constructs and generation of mouse CART19

Plasmids encoding the CAR construct in the SFG γ-retroviral vector [79] were used to transfect gpg29 fibroblasts (H29) with the ProFection Mammalian Transfection System (Promega, WI, USA) according to manufacturer’s instructions in order to generate vesicular stomatitis virus G-glycoprotein-pseudotyped retroviral supernatants. These retroviral supernatants were used to construct stable Moloney murine leukemia virus-pseudotyped retroviral particle-producing Phoenix-ECO cell lines. The SFG-m1928z vector was constructed by stepwise Gibson Assembly (New England BioLabs, MA, USA) using the cDNA of previously described anti-mouse CD19 scFv [31], Myc-tag sequence (EQKLISEEDL), murine CD28 transmembrane and intracellular domain, murine CD3ζ intracellular domain without the stop codon, and P2A self-cleaving peptide.

Splenocytes were harvested from euthanized BALB/c mice in a sterile fashion and processed using a 40 μm nylon cell strainer using complete media (preparation described above). Erythrocytes were depleted using ACK lysing buffer. Using the EasySep Mouse T Cell Isolation Kit (StemCell, Vancouver, Canada), CD3^+^T cells were isolated from the splenocyte pool via negative selection and confirmed as such using flow cytometry. The mouse CD3^+^T cells were then activated using anti CD3/CD28 Dynabeads (Gibco, Thermo Fischer Scientific, #01279757) and expanded in RPMI-1640 medium, supplemented with 10% FBS (Benchmark, #100-106-500), 1% penicillin-streptomycin (Life Technologies, #15140-122), and human IL-2 (50 IU/ml). T cells were then spinoculated with the retroviral supernatant collected from Phoenix-ECO cells at RCF 800g and 32°C, for 35 min and incubated overnight in Retronectin-coated 6-well plates (Lonza, Takara Bio Cat. T100A). After second spinoculation, T cells were assessed using flow cytometry for evaluation of T cells transduction and were rested overnight before intravenous injection into mice.

### *In vitro* Irradiation

Target human and mouse lymphoma cells were cultured at a density of 1 × 10^6^ cells/mI in Iscove’s Modified Dulbecco’s Medium (IMDM, Thermo Fisher Scientific, #12440-061) medium or in Roswell Park Memorial Institute (RPMI-1640, Thermo Fischer Scientific, #11875119) medium, supplemented with 10% fetal bovine serum (FBS, Benchmark, #100-106-500) and 1% penicillin-streptomycin (Life Technologies, #15140-122) then plated in 200-500 µL aliquots into 24- or 48-well plates. After resting for 2 hours in standard culture conditions, cells were exposed to a single fraction of 1-2 Gy irradiation using an Xstrahl Small Animal Radiation Research Platform irradiator (SARRP; Xstrahl, Georgia, United States).

### Flow cytometry

Human cells were stained with different fluorochrome-conjugated monoclonal antibodies specific for CD19, FAS, TRAIL-R1, and TRAIL-R2 **(Supplementary Table 3)** with 4′,6- diamidino-2-phenylindole (DAPI, Life Technologies, CA, USA) as viability dyes. Antigen density of CD19 at 24 h in unirradiated and irradiated cells at 1Gy was quantified by measuring antigen binding capacity (ABC) using BD Quantibrite^TM^ PE beads. Flow cytometry was performed on a BD LSR-II (using high-throuput screening for absolute count) or BD LSRFortessa™ Cell Analyzer. Data were analyzed with FlowJo 10.8.1 (BD Bioscience).

Mouse cells were stained with different fluorochrome-conjugated monoclonal mouse-specific antibodies for CD45, CD3, CD19, CD4, CD8, Myc, Fas (**Supplementary Table 3**). DAPI (Life Technologies, CA, USA) was used as a viability stain. Countbright beads (Invitrogen, MA, USA) were used to determine the absolute number of cells for the *in vivo* studies. Data were acquired on a BD LSR-II or BD LSRFortessa™ Cell Analyzer and analyzed in FlowJo v.10 (BD).

### *In vitro* cytotoxicity assays

Human CART19 cells and control T cells from the same donor (effector cells) were thawed on the day of the experiment. Mouse CART19 cells were manufactured as described above. Human target cells were labeled with cell trace carboxyfluorescein succinimidyl ester (CFSE; life technologies #34554) and plated into 96-well plates at a concentration of 200,000 cells/mL. Mouse target cells (A20 GFP-LUC) expressing GFP were directly plated into 96-well plates at a concentration of 200,000 cells/mL. Target cells were irradiated at 1Gy (or 0Gy for mock radiation control) and then co-cultured with effector cells at different effector: target (E: T) ratios or without effector cells as control. The viability of CFSE-stained cells (human) or GFP^+^ (mouse) was evaluated by flow cytometry using a DAPI stain (Life Technologies, CA, USA). Cytotoxicity was calculated as the absolute number of living target cells in the presence of effector cells relative to the number of control cells cultured without effector cells. Viability was expressed by normalizing to control target cells receiving the same amount of radiation*, i.e.* 0Gy *(mock)* or 1Gy. Cytotoxicity=1-[number of live target cells÷number of live target cells in the control]x100.

### *In vivo* experiments

All experiments were approved by Weill Cornell Medicine Institutional Animal Care & Use Committee (IACUC, #2010-0068). BALB/c mice were purchased from Charles River Laboratories. All mice used were between 6 and 10 weeks old and kept in a pathogen-free facility. Mice were intravenously injected in the tail vein with 5 x10^6^ A20 GFP-LUC cells. Mice were imaged at day 13 with a bioluminescent imager using an IVIS Spectrum In Vivo Imaging System (Perkin Elmer, Connecticut, US) 10 minutes after 200 uL intraperitoneal injections of reconstituted VivoGlo Luciferin (15mg/ml, Promega, Cat. P1043) to confirm engraftment, as we previously described [80]. They were then separated into treatment groups. Cyclophosphamide groups received 100 mg/kg intraperitoneally of reconstituted cyclophosphamide diluted at a 10 mg/mL concentration. The radiation groups received 1Gy of TBI. The mouse CD19 CAR T groups received 1.6 million mouse anti-CD19 CAR T through the tail vein. Mice were monitored weekly with the IVIS imaging system (IVIS spectrum, Caliper LifeSciences, Massachusetts, United States), and imaging analysis was performed via the Living Image Software^®^. Per IACUC guidelines, mice were euthanized when moribund or upon exhibiting significant weight loss, hind limb paralysis, or 3 out of 4 of the following features: ill appearance, weakness, defensive posture, and ruffled fur. The liver, spleen, and bone marrow were collected upon euthanasia. Organs were then processed using 40uM Nylon (Corning, Arizona, United States) cell strainers to create cell suspensions. Erythrocytes were lysed using ACK lysing buffer and cells were stained and run for flow cytometry or sorted for single RNA sequencing.

### A20 rechallenge

Long-term surviving BALB/c mice treated with mCART19 cells and TBI(1Gy) were rechallenged 260 days after the initial tumor injection and rechallenged with additional 5x10^6^ A20 GFP-LUC cells intravenously. They were monitored with the above-mentioned imaging, and peripheral blood draws post-rechallenge. Survival was monitored over time.

### Cytokine analysis

Submandibular peripheral blood was obtained from BALB/c mice and 25-50 uL of whole blood was collected, on day 3 and 9 post-mCART19 administration. Plasma was separated from whole blood by centrifugation at 2700 rpm for 7 minutes then stored at - 80. Samples were thawed, and cytokines were quantified using LEGENDplex™ MU Cytokine Release Syndrome Panel (15-plex) w/VbP by flow cytometry. Data were analyzed with the LEGENDplex™ Data Analysis Software (BioLegend, CA, USA).

### *In vivo* assessment of lymphodepletion

BALB/c mice as described above were separated into four treatment groups: healthy mice (n=3), cyclophosphamide (Cy) only (n=3), Cy + TBI(1Gy) (n=3) and TBI(1Gy) only (n=3). Mice treated with cyclophosphamide received intraperitoneally 100 mg/kg of reconstituted cyclophosphamide diluted at a 10 mg/mL concentration based on individual body weight. Mice in radiation groups received 1Gy of TBI (Rad Source Technologies RS 2000 Biological Research X-ray Irradiator). Mice were then monitored with submandibular peripheral blood draws (40 uL) post treatment at days 3, 12, and 60. Whole blood was run using the Heska HemaTrue Veterinary Hematology Analyzer (Heska, Colorado, US) to assess for white blood cell count, lymphocyte count, neutrophil count, platelets, and hemoglobin. Remaining blood was processed and prepared for flow cytometry as described above.

### *In vivo* assessment of the effect of cyclophosphamide

BALB/c mice were separated into two treatment groups, untreated (n=2) and cyclophosphamide only (n=4). All animals were injected with 5X10^6^ A20 GFP-LUC cells and tumor was allowed to inoculate for 14 days. Tumor progression was monitored through intraperitoneal injections of reconstituted VivoGlo Luciferin (Promega, Cat. P1043) and the IVIS Spectrum In Vivo Imaging System (Perkin Elmer). Animals were randomized based on tumor radiance from imaging and the cyclophosphamide group received intraperitoneally 100 mg/kg of reconstituted cyclophosphamide diluted at a 10 mg/mL concentration based on individual body weight. Mice were monitored weekly per above imaging system until they all succumbed to disease.

### *In vivo* assessment of anti-CD19 CAR T migration

BALB/c mice were injected with 5X10^6^ A20 GFP-LUC as described above and separated into five groups: untreated (n=3), Cy + TBI(1Gy) (n=3), Cy + mCART19 (n=4), Cy + TBI(1Gy) + mCART19 (n=4), Cy + TBI(1Gy) + mCART19 + additional dose of radiation (n=4). Mice were euthanized on Days 3, and 9, organs were harvested to assess infiltration of CART19 to different liver and spleen.

### Statistical analysis

All comparisons between two groups were performed using GraphPad Prism software (GraphPad). Statistical analysis is indicated in the figure legend. Significance was assumed with ^∗^p < 0.05; ^∗∗^p < 0.01; ^∗∗∗^p < 0.001; ^∗∗∗∗^p < 0.0001.

### Single cell RNA sequencing and analysis

#### Sample preparation

Single-cell RNA-seq libraries were prepared according to 10x Genomics specifications (Single Cell 3′ Reagent Kits v3.1 User Guide CG000204, 10x Genomics, Pleasanton, CA, USA). Cellular suspensions (75% viable) at a concentration between 600-1000 cells/µl, were loaded onto to the Chromium Controller to generate barcoded single-cell GEMs, targeting about 10000 single cells per sample. GEM-Reverse Transcription (53 °C for 45 min, 85 °C for 5 min; held at 4 °C) was performed in a C1000 Touch Thermal cycler with 96-Deep Well Reaction Module (Bio-Rad, Hercules). After RT reaction, GEMs were broken, and the single-strand cDNA was cleaned up with DynaBeads MyOne Silane Beads (Thermo Fisher Scientific, Waltham, MA). The cDNA was amplified for 11 cycles (98 °C for 3 min; 98 °C for 15 s, 63°C for 20 s, 72 °C for 1). Quality of the cDNA was assessed using an Agilent Bioanalyzer 2100 (Santa Clara, CA), obtaining a product of about 1100bp. This cDNA was enzymatically fragmented, end repaired, A-tailed, subjected to a double-sided size selection with SPRIselect beads (Beckman Coulter, Indianapolis, IN) and ligated to adaptors provided in the kit. A unique sample index for each library was introduced through 14 cycles of PCR amplification using the indexes provided in the kit (98 °C for 45 s; 98 °C for 20 s, 54 °C for 30 s, and 72 °C for 20 s x 14 cycles; 72 °C for 1 min; held at 4 °C). Indexed libraries were subjected a second double-sided size selection, and libraries were then quantified using Qubit fluorometric quantification (Thermo Fisher Scientific, Waltham, MA). The quality was assessed on an Agilent Bioanalyzer 2100, obtaining an average library size of 470bp.

#### Sequencing and post processing of data

Libraries were diluted to 10nM and clustered on an Illumina NovaSeq 6000 on a pair end read flow cell and sequenced for 28 cycles on R1 (10x barcode and the UMIs), followed by 10 cycles of I7 Index (sample Index), and 91 bases on R2 (transcript), with a coverage around 250M reads per sample. Primary processing of sequencing images was done Illumina’s Real Time Analysis software (RTA).

#### scRNA-seq bioinformatics

scRNA-seq data from the 6 mice were mapped to mm10 mouse genome reference using 10x Genomics Cell Ranger v7.2.0 [81]. Scanpy package [82] was used to create AnnData objects for each of the samples keeping only cells with more than 200 expressed genes and genes with more than 3 counts. In a posterior filtering, only cells with less than 3,000 and more than 50 total counts, genes with more than 50 counts and cells with less than 8% of mitochondrial genes were kept. Doublet estimation was run using Scrublet from scanpy external tools with default parameters [83]. All samples were then concatenated in a unique object and normalization and log transformation steps were run using *sc.pp.normalize_total* and *sc.pp.log1p* followed by scaling data and PCA by *sc.pp.scale* and *sc.tl.pca* functions from scanpy. To correct for sampling batch effects, Harmony integration approach [84] was used by scanpy external tools (*sc.external.pp.harmony_integrate*) and using default parameters. After integration, the nearest neighbors distance matrix was computed using PCA values estimated by harmony and by *sc.pp.neighbors*. Then cells were clustered by Louvain approach [85] and embedding was run through UMAP [86] (*sc.tl.louvain*, *sc.tl.umap*). Marker detection for each identified cluster was run by sc.tl.rank_genes_groups using “Wilcoxon rank-sum” test and general cell type identification was run using over representation analysis approach by decoupler package [87] and using PanglaoDB Immune System database with only canonical markers [88] and by top markers.

Once main cell types were identified, T cells were subtracted and an additional batch correction and reclustering was performed using default parameters by running *sc.external.pp.harmony_integrate*, *sc.pp.neighbors* using harmony’s PCA, *sc.tl.louvain* and *sc.tl.umap*. Annotation of subset of T cells was done based on a combinatorial selection of top markers (sc.tl.rank_genes_groups) and known markers. The two clusters of Naïve T cells (N_1, N_2) were identified by *Lef1and Bach2* expression, one cluster of CD8 expressing T cells (CD8) was identified by *Cd8a*, *Nkg7* and *Ly6c2*, one T effector cluster (EFF) was defined by *Cd44* and *Id2*, and 2 effector clusters showing cytotoxicity (eCTL_1, eCTL_2) were defined by *Nkg7*, *Lgals1*, *Pclaf* and *Stmn1*. Two T regulatory clusters (Treg_1, Treg_2) were defined by high expression of *Il2ra, Ctla4*, *Foxp3* and *Ikzf2*, T follicular helper (T_FH) cluster was identified by *Pifo* and *Gm42722*, T helper type 1 (T_H) cluster was identified by *Iigp1* and *Slfn1* and a cluster showing ribosomal genes as top markers was addressed as T_ribosomal (T_RIBO).

Changes on the proportion of T cells and their subsets between groups were calculated based on compositional data analysis approach. For this purpose, scCODA package was used with a false discovery ratio below 0.15 for main cell types and 0.05 for T cell subsets [89]. For T cell subsets, differential gene expression analysis (DEA) between the two conditions was performed using a pseudocount approach. Using decoupler package pseudocount samples were created summing counts from gene and cells per sample. DESeq2 [90] package was used to run DEA between conditions on cell type specific bases and genes with padj < 0.05 and absolute log fold change of 0.25 were called “dysregulated”. Gene Set Enrichment Analysis was performed using *get_gsea_df* function from decoupler based on GSEA software [91, 92], and ranking genes in cell type specific manner using the Wald statistic (stat) obtained from DEA by DESeq2. For this step mouse specific hallmark gene sets were used [53]. To assess synthetic mCART19 retention and expansion within our T cell clusters, we defined mCART19 positive cells as those expressing at least one raw count of Myc gene. Next, we assessed the enrichment of Myc-positive (synthetic mCART19) cells between our conditions using a cell type-specific binomial test and under FDR padj < 0.05. This approach allowed us to estimate whether the likelihood of finding synthetic CAR T cells in any of our conditions was significantly higher or lower than expected by chance. Then we run pseudocount DEA between conditions using all cells identified as mCART19 positive cells irrelevantly of the subsets identified before. Heatmaps were constructed using z-score values of pseudobulk gene expression at the level of the replicate; z-scores were calculated per gene and all values were normalized to a range of 0-1 Using the R packages CompleHeatmap and ggplot2.

All the code used for scRNA-seq analysis will be available at (https://github.com/abcwcm/alhomoud_cart_rad).

## AUTHORS’ CONTRIBUTIONS

**MA:** experimentation, data analysis, validation, methodology, writing original draft, writing review, and editing. **MF:** experimentation, data analysis, validation, methodology, writing original draft, writing review, and editing. **MS:** data generation, methodology. **JAF:** analysis, methodology, validation. **SY:** methodology, writing review, and editing. **LM:** data generation. **KR:** critical review of the manuscript, feedback on CRS-like scoring. **MA:** formal analysis, methodology, validation. **DB:** formal analysis, methodology, validation. **RB:** resources, writing review, and editing. **KVB:** supervision, writing review, and editing. **LG:** resources, methodology, writing review, and editing. **OB:** resources, supervision, writing review, and editing. **JM:** data generation, analysis, supervision, writing review, and editing. **SF:** conceptualization, resources, supervision, funding acquisition, methodology, writing review, and editing. **MLG:** conceptualization, resources, supervision, funding acquisition, project administration, validation, writing, and editing.

## ACKNOWLEDGMENTS

The authors thank Alicia Alonso (genomics cores-WCMC) and Sharmaine Griffith-Baker (Biorepository and personalized medical center-WCMC) for their cell sorting and single-cell RNA sequencing expertise. We thank Alexandra Gomez-Arteaga, Roni Shouval, Paul Pagnini, Tsiporah Shore, and Juliet N Barker for the thoughtful comments and feedback on the manuscript. This project is supported in part by MCC Clinical Trials Innovation Fund to MLG, SY, LG and SF. Grant/Research support from SF: Bristol Myers Squibb, Varian, Regeneron, Merck, Celldex, Arcus, DoD - BCRP (#BC180476, #BC180595, BC201085P3), NIH U54 CA 27429. Grant/Research support from OB, JM: Foundation Charles Nicolle, the Canceropole Nord-Ouest, the GCS/G4” and the Normandie region. Grant/Research support from MLG: R01 CA098571, R01 CA080728.

## SUPPLEMENTARY FIGURE LEGENDS

**Supplementary Figure 1. Gating strategies for assessment of DRs, and cytotoxicity. A.** Gating strategy for assessment of FAS and TRAIL-R2, when using flow cytometry (representative example; OCI-LY10). Representative histograms are shown for cells irradiated with 0Gy (mock) and 1Gy overlaying with unstained control. **B.** Fold change of MFI for FAS and TRAIL-R2 in OCI-LY10 measured by flow cytometry over multiple time points after radiation (1Gy); 3, 6, 9, and 24 hours. **C.** Gating strategy used for the cytotoxicity assays; cells were gated as living CFSE^+^ cells for targets cells.

**Supplementary Figure 2. Manufacturing of mCART19, gating strategies for the mCART19 purity and cytotoxicity against A20 assessments. A.** Schematic diagram of mCART19 production. T cells were isolated from spleens of healthy BALB/c mice, activated (anti-CD3/CD28 beads), and transduced with SFG-m1928z vector. **B.** Gating strategy for assessing mouse T cells purity gated as CD3^+^ cells. **C**. Gating strategy for assessing mCART19 cells gated as CD3^+^/Myc^+^ cells. **D.** Gating strategy for the cytotoxicity assay; GFP^+^ cells (A20) were gated for further evaluation of viability using DAPI.

**Supplementary Figure 3. The lymphodepletion regimen does not eliminate LBCL mouse model. A. Left,** Schematic representation of the experimental setup for Cy conditioning in the A20 lymphoma mouse model. BALB/c mice injected with A20 GFP-LUC. Lymphoma burden was assessed using BLI. Cohorts were assigned at day 13 as follows 1: saline control (n= 2), 2: Cy at 100mg/kg (n=4). **Right.** Lymphoma progression represented by average radiance (p/sec/cm2/sr) for the treatment groups at the indicated times. Each circle represents the mean for the group, error is indicated by SD. **B**. Gating strategy for assessing mouse B cells in peripheral blood gated as CD45^+^/CD19^+^GFP^-^cells.

**Supplementary Figure 4. Assessment of the effects of TBI(1Gy) on mCART19 expansion and cytokine release. A.** Graphical representation for the experimental setup describing lymphodepletion and long-term cytopenia assessment in immunocompetent mouse model. BALB/c mice were treated with different conditioning regimens as indicated; 1: Saline control (n=3); 2: Cy (n=3); 3: TBI(1Gy) (n=3); 4: Cy/TBI(1Gy) (n=3). Complete blood count was monitored using peripheral blood draws**. B.** Schematic diagram describing assessment of mCART19 expansion in peripheral blood of mice. BALB/c mice were I.V. injected with A20 GFP-LUC and engraftment was confirmed by BLI 13 days after injection. Cohorts were as indicated: 1: Saline control (n=5); 2:Cy/TBI(1Gy) (n=5); 3: Cy/mCART19 (n=5); 4: Cy/TBI(1Gy)/mCART19 (n=5). TBI(1Gy) was given 4 hours before mCART19 injection. Peripheral blood draws were performed on day 9 to assess for mCART19 by flow cytometry. **C.** Gating strategy for evaluation mCART19 in peripheral blood gated as CD3^+^Myc^+^ cells. **D.** Heatmap representing the normalized Z-score for cytokine concentrations in the peripheral blood of mice on Day 3 and Day 9 post-treatment. Each cell represents an average of the animals in the cohort evaluated in duplicates (n=5). **E.** Cytokine release-like syndrome clinical manifestation scoring (described in supplementary table 1) of mice receiving Cy/mCART19 vs. Cy/radiation/mCART19. The clinical score measured at day -1 (pre-treatment) and day +3 (post-treatment). *P*-value calculated using two-way ANOVA.

**Supplementary Figure 5. A.** Percent of mCART19 in bone marrow of mice sacrificed at 62 days post rechallenge from Fig 4(D-F), is shown, each dot represents one mouse, and the bar represents the mean with SD. *P*-value calculated using unpaired t-test. **B.** BLI tumor growth measured over time for the rechallenged mice represented by average radiance (p/sec/cm2/sr).

**Supplementary Figure 6. A. Upper.** Volcano plot of Wilcoxon rank-sum scores from the differential gene expression analysis of Cy/TBI(1Gy)/mCART19 vs Cy/mCART19 in Naïve_1 T cell subset. Genes with adjusted *P*-value=1 were filtered out before plotting. **Lower.** Top 10 enriched pathways on Naïve_1 T cell subset in Cy/TBI(1Gy)/mCART19 vs. Cy/mCART19. **B.** Dot plot showing main gene markers used to classify general cell types**. C.** Dot plot showing top markers genes used to classify T cell subsets.

## REFERENCES

1. Neelapu SS, Locke FL, Bartlett NL, Lekakis LJ, Miklos DB, Jacobson CA, et al. Axicabtagene Ciloleucel CAR T-Cell Therapy in Refractory Large B-Cell Lymphoma. N Engl J Med. 2017;377(26):2531–44.

2. Abramson JS, Palomba ML, Gordon LI, Lunning MA, Wang M, Arnason J, et al. Lisocabtagene maraleucel for patients with relapsed or refractory large B-cell lymphomas (TRANSCEND NHL 001): a multicentre seamless design study. Lancet. 2020;396(10254):839–52.

3. Schuster SJ, Bishop MR, Tam CS, Waller EK, Borchmann P, McGuirk JP, et al. Tisagenlecleucel in Adult Relapsed or Refractory Diffuse Large B-Cell Lymphoma. N Engl J Med. 2019;380(1):45–56.

4. Neelapu SS, Jacobson CA, Ghobadi A, Miklos DB, Lekakis LJ, Oluwole OO, et al. Five-year follow-up of ZUMA-1 supports the curative potential of axicabtagene ciloleucel in refractory large B-cell lymphoma. Blood. 2023;141(19):2307–15.

5. Jain MD, Spiegel JY, Nastoupil LJ, Tamaresis J, Ghobadi A, Lin Y, et al. Five-Year Follow-Up of Standard-of-Care Axicabtagene Ciloleucel for Large B-Cell Lymphoma: Results From the US Lymphoma CAR T Consortium. J Clin Oncol. 2024:Jco2302786.

6. Schaue Dr, Comin-Anduix B, Ribas A, Zhang L, Goodglick L, Sayre JW, et al. T-Cell Responses to Survivin in Cancer Patients Undergoing Radiation Therapy. Clinical Cancer Research. 2008;14(15):4883–90.

7. Burnette BC, Liang H, Lee Y, Chlewicki L, Khodarev NN, Weichselbaum RR, et al. The efficacy of radiotherapy relies upon induction of type i interferon-dependent innate and adaptive immunity. Cancer Res. 2011;71(7):2488–96.

8. Lee Y, Auh SL, Wang Y, Burnette B, Wang Y, Meng Y, et al. Therapeutic effects of ablative radiation on local tumor require CD8+ T cells: changing strategies for cancer treatment. Blood. 2009;114(3):589–95.

9. Golden EB, Frances D, Pellicciotta I, Demaria S, Helen Barcellos-Hoff M, Formenti SC. Radiation fosters dose-dependent and chemotherapy-induced immunogenic cell death. Oncoimmunology. 2014;3:e28518.

10. Deng L, Liang H, Xu M, Yang X, Burnette B, Arina A, et al. STING-Dependent Cytosolic DNA Sensing Promotes Radiation-Induced Type I Interferon-Dependent Antitumor Immunity in Immunogenic Tumors. Immunity. 2014;41(5):843–52.

11. Shaverdian N, Lisberg AE, Bornazyan K, Veruttipong D, Goldman JW, Formenti SC, et al. Previous radiotherapy and the clinical activity and toxicity of pembrolizumab in the treatment of non-small-cell lung cancer: a secondary analysis of the KEYNOTE-001 phase 1 trial. The lancet oncology. 2017;18(7):895–903.

12. Kline C, Liu SJ, Duriseti S, Banerjee A, Nicolaides T, Raber S, et al. Reirradiation and PD-1 inhibition with nivolumab for the treatment of recurrent diffuse intrinsic pontine glioma: a single-institution experience. Journal of neuro-oncology. 2018;140:629–38.

13. Luke JJ, Lemons JM, Karrison TG, Pitroda SP, Melotek JM, Zha Y, et al. Safety and clinical activity of pembrolizumab and multisite stereotactic body radiotherapy in patients with advanced solid tumors. Journal of Clinical Oncology. 2018;36(16):1611–8.

14. Koller KM, Mackley HB, Liu J, Wagner H, Talamo G, Schell TD, et al. Improved survival and complete response rates in patients with advanced melanoma treated with concurrent ipilimumab and radiotherapy versus ipilimumab alone. Cancer biology & therapy. 2017;18(1):36–42.

15. Galluzzi L, Aryankalayil MJ, Coleman CN, Formenti SC. Emerging evidence for adapting radiotherapy to immunotherapy. Nat Rev Clin Oncol. 2023;20(8):543–57.

16. DeSelm C, Palomba ML, Yahalom J, Hamieh M, Eyquem J, Rajasekhar VK, et al. Low-Dose Radiation Conditioning Enables CAR T Cells to Mitigate Antigen Escape. Mol Ther. 2018;26(11):2542–52.

17. Sugita M, Yamazaki T, Alhomoud M, Martinet J, Latouche JB, Golden E, et al. Radiation therapy improves CAR T cell activity in acute lymphoblastic leukemia. Cell Death Dis. 2023;14(5):305.

18. Sim AJ, Jain MD, Figura NB, Chavez JC, Shah BD, Khimani F, et al. Radiation Therapy as a Bridging Strategy for CAR T Cell Therapy With Axicabtagene Ciloleucel in Diffuse Large B-Cell Lymphoma. Int J Radiat Oncol Biol Phys. 2019;105(5):1012–21.

19. Pinnix CC, Gunther JR, Dabaja BS, Strati P, Fang P, Hawkins MC, et al. Bridging therapy prior to axicabtagene ciloleucel for relapsed/refractory large B-cell lymphoma. Blood Adv. 2020;4(13):2871–83.

20. Christopher W, al e. Bridging Radiation Therapy Before Commercial Chimeric Antigen Receptor T-Cell Therapy for Relapsed or Refractory Aggressive B-Cell Lymphoma. 2020.

21. Claire R, al e. Effective bridging therapy can improve CD19 CAR-T outcomes while maintaining safety in patients with large B-cell lymphoma. Blood Advances. 2023.

22. Colton L, al e. Long-Term Follow-Up of Bridging Therapies Prior to CAR T-Cell Therapy for Relapsed/Refractory Large B Cell Lymphoma. Cancers. 2023.

23. Deshpande A, Rule W, Rosenthal A. Radiation and Chimeric Antigen Receptor T-cell Therapy in B-cell Non-Hodgkin Lymphomas. Current Treatment Options in Oncology. 2022.

24. Forat L, al e. The impact of bridging therapy prior to CD19-directed chimeric antigen receptor T-cell therapy in patients with large B-cell lymphoma. 2021.

25. Harper H, al e. Bridging Radiation Rapidly and Effectively Cytoreduces High-Risk Relapsed/Refractory Aggressive B Cell Lymphomas Prior to Chimeric Antigen Receptor T Cell Therapy. Transplant Cell Therapy. 2023.

26. Omran S, al e. Don’t Put the CART Before the Horse: The Role of Radiation Therapy in Peri-CAR T-cell Therapy for Aggressive B-cell Non-Hodgkin Lymphoma. International Journal of Radiation Oncology. 2022.

27. Alhomoud M, Ibrahim R, Chen Z, Martinet J, Foley M, Gomez-Arteaga A, et al. Safety and Efficacy of Bridging Radiation Therapy in the Context of CD19 CART Therapy for Non-Hodgkin Lymphoma: Systematic Review and Meta-Analysis. Transplantation and Cellular Therapy, Official Publication of the American Society for Transplantation and Cellular Therapy. 2024;30(2):S228.

28. Locke FL, Filosto S, Chou J, Vardhanabhuti S, Perbost R, Dreger P, et al. Impact of tumor microenvironment on efficacy of anti-CD19 CAR T cell therapy or chemotherapy and transplant in large B cell lymphoma. Nature Medicine. 2024;30(2):507–18.

29. Eugene-Norbert M, Cuffel A, Riou G, Jean L, Blondel C, Dehayes J, et al. Development of optimized cytotoxicity assays for assessing the antitumor potential of CAR-T cells. J Immunol Methods. 2024;525:113603.

30. Kuhn NF, Purdon TJ, van Leeuwen DG, Lopez AV, Curran KJ, Daniyan AF, et al. CD40 Ligand-Modified Chimeric Antigen Receptor T Cells Enhance Antitumor Function by Eliciting an Endogenous Antitumor Response. Cancer Cell. 2019;35(3):473–88.e6.

31. Davila ML, Kloss CC, Gunset G, Sadelain M. CD19 CAR-targeted T cells induce long-term remission and B Cell Aplasia in an immunocompetent mouse model of B cell acute lymphoblastic leukemia. PLoS One. 2013;8(4):e61338.

32. Turtle CJ, Hanafi LA, Berger C, Hudecek M, Pender B, Robinson E, et al. Immunotherapy of non-Hodgkin’s lymphoma with a defined ratio of CD8+ and CD4+ CD19-specific chimeric antigen receptor-modified T cells. Sci Transl Med. 2016;8(355):355ra116.

33. Fischer L, Grieb N, Born P, Weiss R, Seiffert S, Boldt A, et al. Cellular dynamics following CAR T cell therapy are associated with response and toxicity in relapsed/refractory myeloma. Leukemia. 2024;38(2):372–82.

34. Wang M, Munoz J, Goy A, Locke FL, Jacobson CA, Hill BT, et al. Three-Year Follow-Up of KTE-X19 in Patients With Relapsed/Refractory Mantle Cell Lymphoma, Including High-Risk Subgroups, in the ZUMA-2 Study. J Clin Oncol. 2023;41(3):555–67.

35. Cappell KM, Sherry RM, Yang JC, Goff SL, Vanasse DA, McIntyre L, et al. Long-Term Follow-Up of Anti-CD19 Chimeric Antigen Receptor T-Cell Therapy. J Clin Oncol. 2020;38(32):3805–15.

36. Locke FL, Miklos DB, Jacobson CA, Perales MA, Kersten MJ, Oluwole OO, et al. Axicabtagene Ciloleucel as Second-Line Therapy for Large B-Cell Lymphoma. N Engl J Med. 2022;386(7):640–54.

37. Kuziel WA, Morgan SJ, Dawson TC, Griffin S, Smithies O, Ley K, et al. Severe reduction in leukocyte adhesion and monocyte extravasation in mice deficient in CC chemokine receptorl’.2. Proceedings of the National Academy of Sciences. 1997;94(22):12053–8.

38. Boring L, Gosling J, Chensue SW, Kunkel SL, Farese RV, Jr., Broxmeyer HE, et al. Impaired monocyte migration and reduced type 1 (Th1) cytokine responses in C-C chemokine receptor 2 knockout mice. J Clin Invest. 1997;100(10):2552–61.

39. Carr MW, Roth SJ, Luther E, Rose SS, Springer TA. Monocyte chemoattractant protein 1 acts as a T-lymphocyte chemoattractant. Proc Natl Acad Sci U S A. 1994;91(9):3652–6.

40. Brown CE, Vishwanath RP, Aguilar B, Starr R, Najbauer J, Aboody KS, et al. Tumor-derived chemokine MCP-1/CCL2 is sufficient for mediating tumor tropism of adoptively transferred T cells. J Immunol. 2007;179(5):3332–41.

41. Di Stasi A, De Angelis B, Rooney CM, Zhang L, Mahendravada A, Foster AE, et al. T lymphocytes coexpressing CCR4 and a chimeric antigen receptor targeting CD30 have improved homing and antitumor activity in a Hodgkin tumor model. Blood, The Journal of the American Society of Hematology. 2009;113(25):6392–402.

42. Craddock JA, Lu A, Bear A, Pule M, Brenner MK, Rooney CM, et al. Enhanced tumor trafficking of GD2 chimeric antigen receptor T cells by expression of the chemokine receptor CCR2b. Journal of immunotherapy. 2010;33(8):780–8.

43. Moon EK, Carpenito C, Sun J, Wang L-CS, Kapoor V, Predina J, et al. Expression of a functional CCR2 receptor enhances tumor localization and tumor eradication by retargeted human T cells expressing a mesothelin-specific chimeric antibody receptor. Clinical cancer research. 2011;17(14):4719–30.

44. Jamali A, Ho N, Braun A, Adabi E, Thalheimer FB, Buchholz CJ. Early induction of cytokine release syndrome by rapidly generated CAR T cells in preclinical models. EMBO Mol Med. 2024;16(4):784–804.

45. Flurkey K, M. Currer J, Harrison DE. Chapter 20 - Mouse Models in Aging Research. In: Fox JG, Davisson MT, Quimby FW, Barthold SW, Newcomer CE, Smith AL, editors. The Mouse in Biomedical Research (Second Edition). Burlington: Academic Press; 2007. p. 637–72.

46. Preglej T, Ellmeier W. CD4(+) Cytotoxic T cells - Phenotype, Function and Transcriptional Networks Controlling Their Differentiation Pathways. Immunol Lett. 2022;247:27–42.

47. Lin W, Singh V, Springer R, Choonoo G, Gupta N, Patel A, et al. Human CD4 cytotoxic T lymphocytes mediate potent tumor control in humanized immune system mice. Communications Biology. 2023;6(1):447.

48. Oh DY, Fong L. Cytotoxic CD4(+) T cells in cancer: Expanding the immune effector toolbox. Immunity. 2021;54(12):2701–11.

49. López-Cantillo G, Urueña C, Camacho BA, Ramírez-Segura C. CAR-T Cell Performance: How to Improve Their Persistence? Front Immunol. 2022;13:878209.

50. Ahmed R, Gray D. Immunological memory and protective immunity: understanding their relation. Science. 1996;272(5258):54–60.

51. Sommermeyer D, Hudecek M, Kosasih PL, Gogishvili T, Maloney DG, Turtle CJ, et al. Chimeric antigen receptor-modified T cells derived from defined CD8+ and CD4+ subsets confer superior antitumor reactivity in vivo. Leukemia. 2016;30(2):492–500.

52. Arcangeli S, Bove C, Mezzanotte C, Camisa B, Falcone L, Manfredi F, et al. CAR T cell manufacturing from naive/stem memory T lymphocytes enhances antitumor responses while curtailing cytokine release syndrome. J Clin Invest. 2022;132(12).

53. Liberzon A, Birger C, Thorvaldsdóttir H, Ghandi M, Mesirov JP, Tamayo P. The Molecular Signatures Database (MSigDB) hallmark gene set collection. Cell Syst. 2015;1(6):417–25.

54. Veiga-Fernandes H, Rocha B. High expression of active CDK6 in the cytoplasm of CD8 memory cells favors rapid division. Nat Immunol. 2004;5(1):31–7.

55. Jones RG, Bui T, White C, Madesh M, Krawczyk CM, Lindsten T, et al. The proapoptotic factors Bax and Bak regulate T Cell proliferation through control of endoplasmic reticulum Ca(2+) homeostasis. Immunity. 2007;27(2):268–80.

56. Good Z, Spiegel JY, Sahaf B, Malipatlolla MB, Ehlinger ZJ, Kurra S, et al. Post-infusion CAR T(Reg) cells identify patients resistant to CD19-CAR therapy. Nat Med. 2022;28(9):1860–71.

57. Hu C, Li T, Xu Y, Zhang X, Li F, Bai J, et al. CellMarker 2.0: an updated database of manually curated cell markers in human/mouse and web tools based on scRNA-seq data. Nucleic Acids Res. 2023;51(D1):D870–d6.

58. Singh N, Lee YG, Shestova O, Ravikumar P, Hayer KE, Hong SJ, et al. Impaired death receptor signaling in leukemia causes antigen-independent resistance by inducing CAR T-cell dysfunction. Cancer discovery. 2020;10(4):552–67.

59. Upadhyay R, Boiarsky JA, Pantsulaia G, Svensson-Arvelund J, Lin MJ, Wroblewska A, et al. A critical role for Fas-mediated off-target tumor killing in T-cell immunotherapy. Cancer discovery. 2021;11(3):599–613.

60. Scholler N, Perbost R, Locke FL, Jain MD, Turcan S, Danan C, et al. Tumor immune contexture is a determinant of anti-CD19 CAR T cell efficacy in large B cell lymphoma. Nature medicine. 2022;28(9):1872–82.

61. Jain MD, Zhao H, Wang X, Atkins R, Menges M, Reid K, et al. Tumor interferon signaling and suppressive myeloid cells are associated with CAR T-cell failure in large B-cell lymphoma. Blood, The Journal of the American Society of Hematology. 2021;137(19):2621–33.

62. Rossi J, Paczkowski P, Shen Y-W, Morse K, Flynn B, Kaiser A, et al. Preinfusion polyfunctional anti-CD19 chimeric antigen receptor T cells are associated with clinical outcomes in NHL. Blood, The Journal of the American Society of Hematology. 2018;132(8):804–14.

63. Fraietta JA, Lacey SF, Orlando EJ, Pruteanu-Malinici I, Gohil M, Lundh S, et al. Determinants of response and resistance to CD19 chimeric antigen receptor (CAR) T cell therapy of chronic lymphocytic leukemia. Nature medicine. 2018;24(5):563–71.

64. Rejeski K, Jain MD, Smith EL. Mechanisms of Resistance and Treatment of Relapse after CAR T-cell Therapy for Large B-cell Lymphoma and Multiple Myeloma. Transplant Cell Ther. 2023;29(7):418–28.

65. Kim AB, Chou SY, Kang S, Kwon E, Inkman M, Szymanski J, et al. Intrinsic tumor resistance to CAR T cells is a dynamic transcriptional state that is exploitable with low-dose radiation. Blood Adv. 2023;7(18):5396–408.

66. Upadhyay R, Boiarsky JA, Pantsulaia G, Svensson-Arvelund J, Lin MJ, Wroblewska A, et al. A Critical Role for Fas-Mediated Off-Target Tumor Killing in T-cell Immunotherapy. Cancer Discovery. 2021;11(3):599–613.

67. Safwat A. The role of low-dose total body irradiation in treatment of non-Hodgkin’s lymphoma: a new look at an old method. Radiother Oncol. 2000;56(1):1–8.

68. Leimert JT, Corder MP, Tewfik HH, Guthrie R, Maguire LC, Gingrich RD. Total body irradiation and cyclophosphamide, vincristine, prednisone in the treatment of favorable prognosis non-Hodgkin’s lymphomas. Int J Radiat Oncol Biol Phys. 1979;5(9):1479–83.

69. Meerwaldt JH, Carde P, Burgers JM, Monconduit M, Thomas J, Somers R, et al. Low-dose total body irradiation versus combination chemotherapy for lymphomas with follicular growth pattern. Int J Radiat Oncol Biol Phys. 1991;21(5):1167–72.

70. Sermer D, Batlevi C, Palomba ML, Shah G, Lin RJ, Perales M-A, et al. Outcomes in patients with DLBCL treated with commercial CAR T cells compared with alternate therapies. Blood advances. 2020;4(19):4669–78.

71. Dean EA, Mhaskar RS, Lu H, Mousa MS, Krivenko GS, Lazaryan A, et al. High metabolic tumor volume is associated with decreased efficacy of axicabtagene ciloleucel in large B-cell lymphoma. Blood Advances. 2020;4(14):3268–76.

72. Henke E, Nandigama R, Ergün S. Extracellular Matrix in the Tumor Microenvironment and Its Impact on Cancer Therapy. Front Mol Biosci. 2019;6:160.

73. Jameus A, Kennedy AE, Thome C. Hematological Changes Following Low Dose Radiation Therapy and Comparison to Current Standard of Care Cancer Treatments. Dose Response. 2021;19(4):15593258211056196.

74. Rejeski K, Subklewe M, Aljurf M, Bachy E, Balduzzi A, Barba P, et al. Immune effector cell-associated hematotoxicity: EHA/EBMT consensus grading and best practice recommendations. Blood. 2023;142(10):865–77.

75. Rejeski K, Jain MD, Shah NN, Perales MA, Subklewe M. Immune effector cell-associated haematotoxicity after CAR T-cell therapy: from mechanism to management. Lancet Haematol. 2024;11(6):e459–e70.

76. Rejeski K, Wang Y, Hansen DK, Iacoboni G, Bachy E, Bansal R, et al. Applying the EHA/EBMT grading for ICAHT after CAR-T: comparative incidence and association with infections and mortality. Blood Adv. 2024;8(8):1857–68.

77. Lemoine J, Bachy E, Cartron G, Beauvais D, Gastinne T, Di Blasi R, et al. Nonrelapse mortality after CAR T-cell therapy for large B-cell lymphoma: a LYSA study from the DESCAR-T registry. Blood Adv. 2023;7(21):6589–98.

78. Cordas Dos Santos DM, Tix T, Shouval R, Gafter-Gvili A, Alberge JB, Cliff ERS, et al. A systematic review and meta-analysis of nonrelapse mortality after CAR T cell therapy. Nat Med. 2024;30(9):2667–78.

79. Rivière I, Brose K, Mulligan RC. Effects of retroviral vector design on expression of human adenosine deaminase in murine bone marrow transplant recipients engrafted with genetically modified cells. Proc Natl Acad Sci U S A. 1995;92(15):6733–7.

80. Alhomoud M, Martinet J, Sugita M, Gomez-Arteaga A, Guzman ML. Chapter 12 - Methods to monitor in vivo expansion and efficacy of CAR-T cells in preclinical models. In: Spada S, Galluzzi L, editors. Methods in Cell Biology. 167: Academic Press; 2022. p. 185–201.

81. Zheng GX, Terry JM, Belgrader P, Ryvkin P, Bent ZW, Wilson R, et al. Massively parallel digital transcriptional profiling of single cells. Nat Commun. 2017;8:14049.

82. Wolf FA, Angerer P, Theis FJ. SCANPY: large-scale single-cell gene expression data analysis. Genome Biology. 2018;19(1):15.

83. Wolf FA, Hamey FK, Plass M, Solana J, Dahlin JS, Göttgens B, et al. PAGA: graph abstraction reconciles clustering with trajectory inference through a topology preserving map of single cells. Genome Biology. 2019;20(1):59.

84. Korsunsky I, Millard N, Fan J, Slowikowski K, Zhang F, Wei K, et al. Fast, sensitive and accurate integration of single-cell data with Harmony. Nature Methods. 2019;16(12):1289–96.

85. Blondel VD, Guillaume J-L, Lambiotte R, Lefebvre E. Fast unfolding of communities in large networks. Journal of Statistical Mechanics: Theory and Experiment. 2008;2008(10):P10008.

86. Becht E, McInnes L, Healy J, Dutertre C-A, Kwok IWH, Ng LG, et al. Dimensionality reduction for visualizing single-cell data using UMAP. Nature Biotechnology. 2019;37(1):38–44.

87. Badia IMP, Vélez Santiago J, Braunger J, Geiss C, Dimitrov D, Müller-Dott S, et al. decoupleR: ensemble of computational methods to infer biological activities from omics data. Bioinform Adv. 2022;2(1):vbac016.

88. Franzén O, Gan L-M, Björkegren JLM. PanglaoDB: a web server for exploration of mouse and human single-cell RNA sequencing data. Database. 2019;2019.

89. Büttner M, Ostner J, Müller CL, Theis FJ, Schubert B. scCODA is a Bayesian model for compositional single-cell data analysis. Nature Communications. 2021;12(1):6876.

90. Love MI, Huber W, Anders S. Moderated estimation of fold change and dispersion for RNA-seq data with DESeq2. Genome Biology. 2014;15(12):550.

91. Subramanian A, Tamayo P, Mootha VK, Mukherjee S, Ebert BL, Gillette MA, et al. Gene set enrichment analysis: a knowledge-based approach for interpreting genome-wide expression profiles. Proc Natl Acad Sci U S A. 2005;102(43):15545–50.

92. Mootha VK, Lindgren CM, Eriksson K-F, Subramanian A, Sihag S, Lehar J, et al. PGC-1α-responsive genes involved in oxidative phosphorylation are coordinately downregulated in human diabetes. Nature Genetics. 2003;34(3):267–73.

