## Supplementary figures and images for "Total body irradiation primes CD19-directed CAR T cells against large B-cell lymphoma"

### supplementary Fig. 1

#### Supplementary Figure 1.

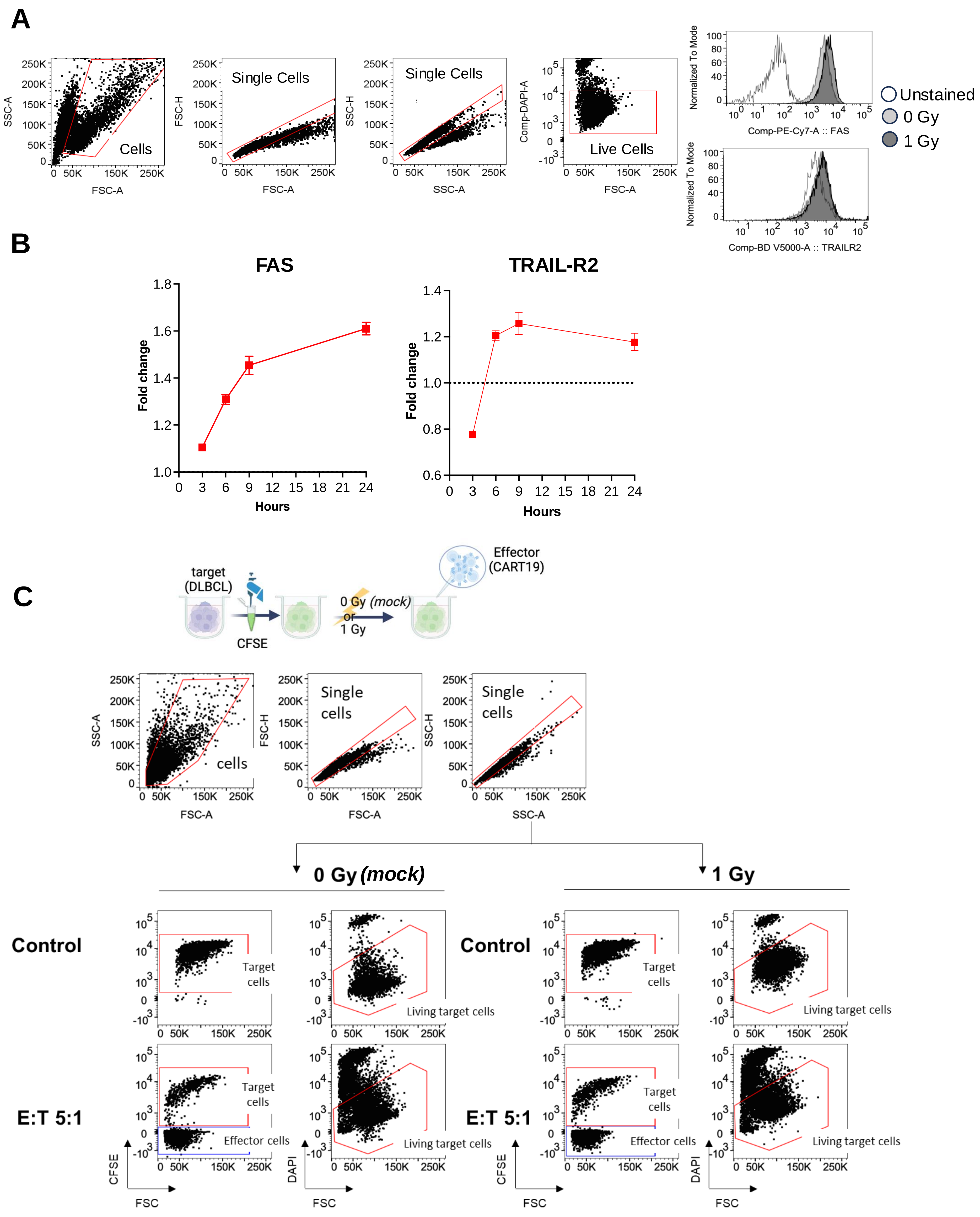

### supplementary Fig. 2

**Supplementary Figure 2.**

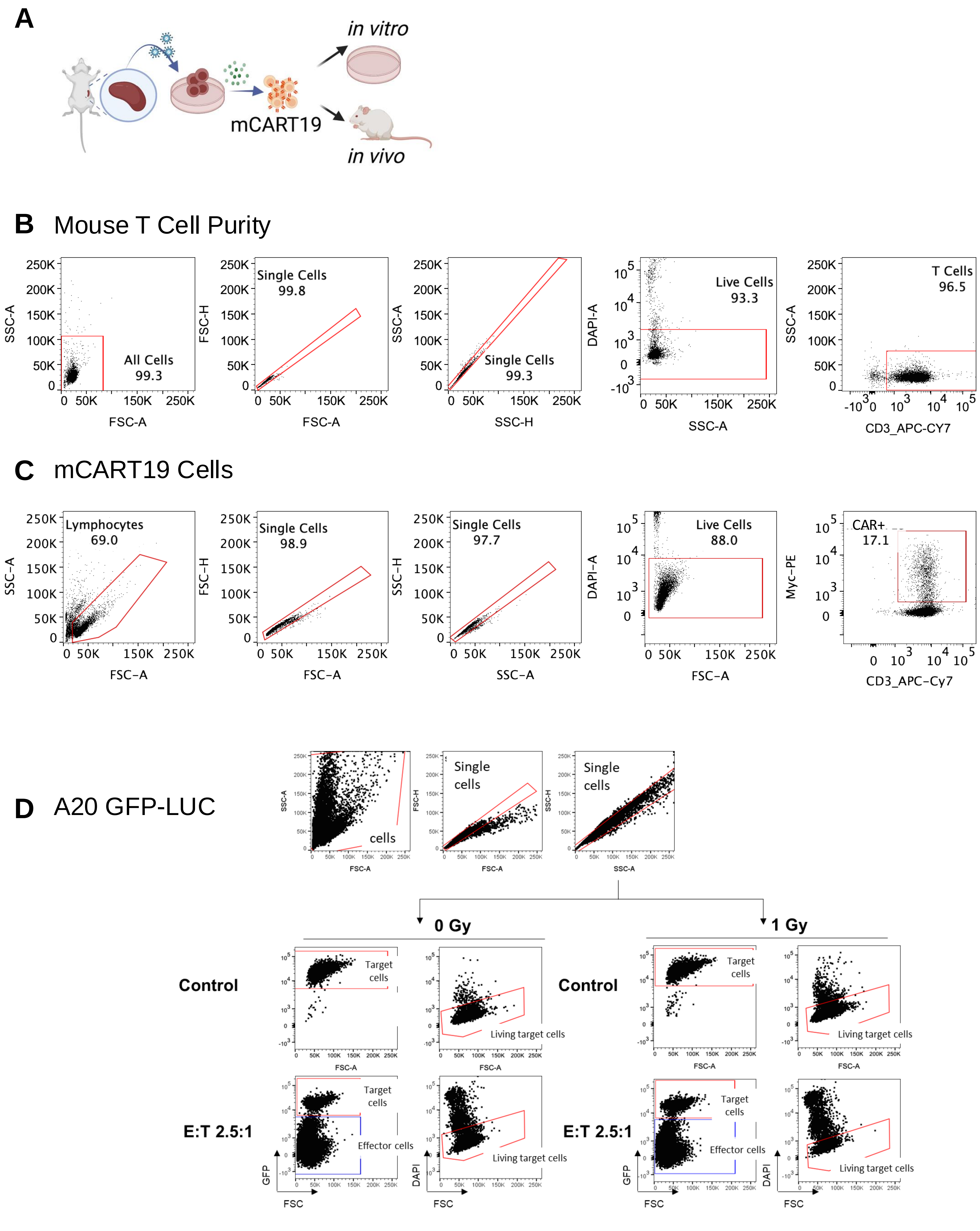

### supplementary Fig. 3

#### Supplementary Figure 3.

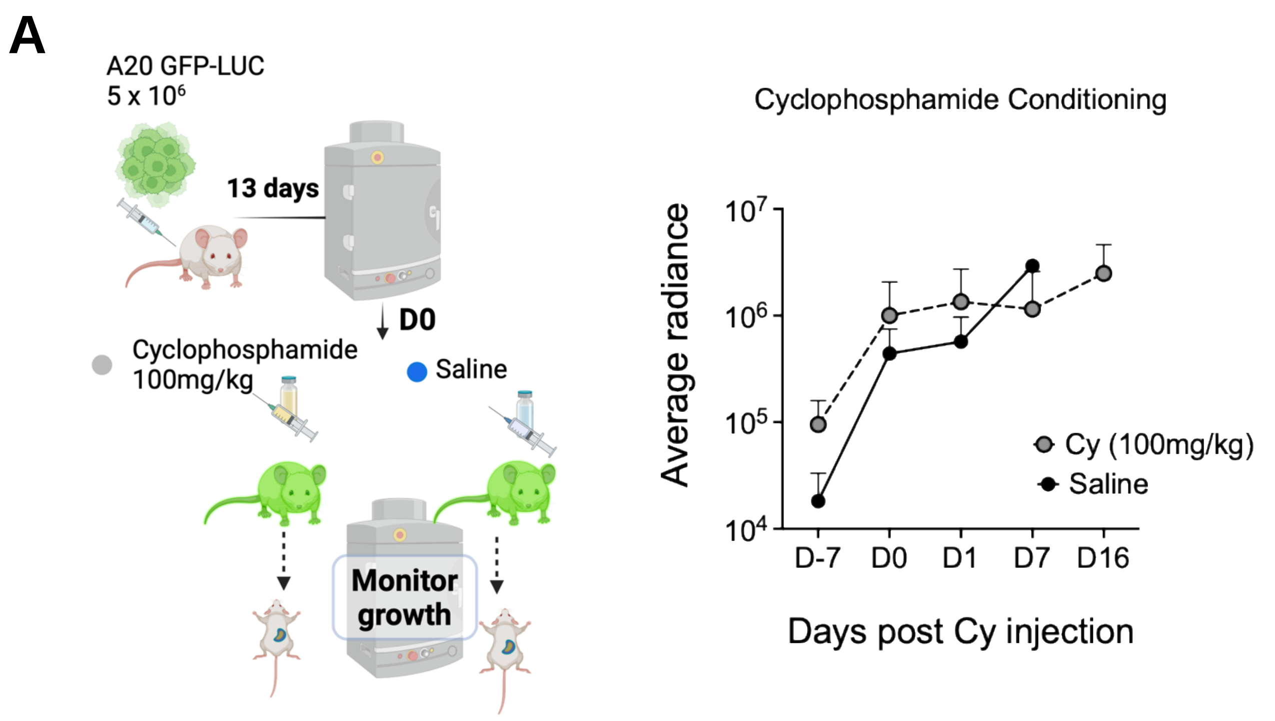


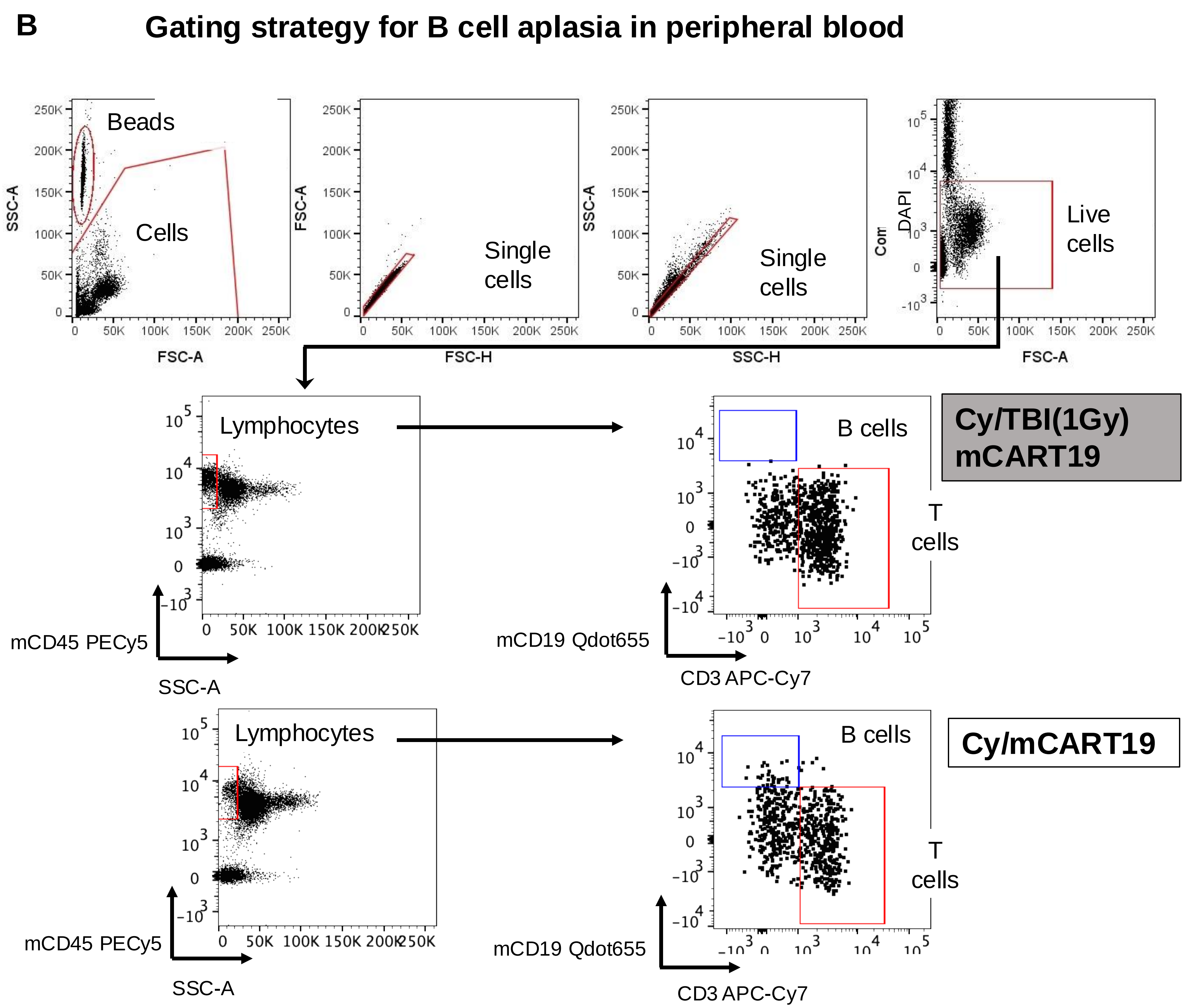

### supplementary Fig. 4

#### Supplementary Figure 4.

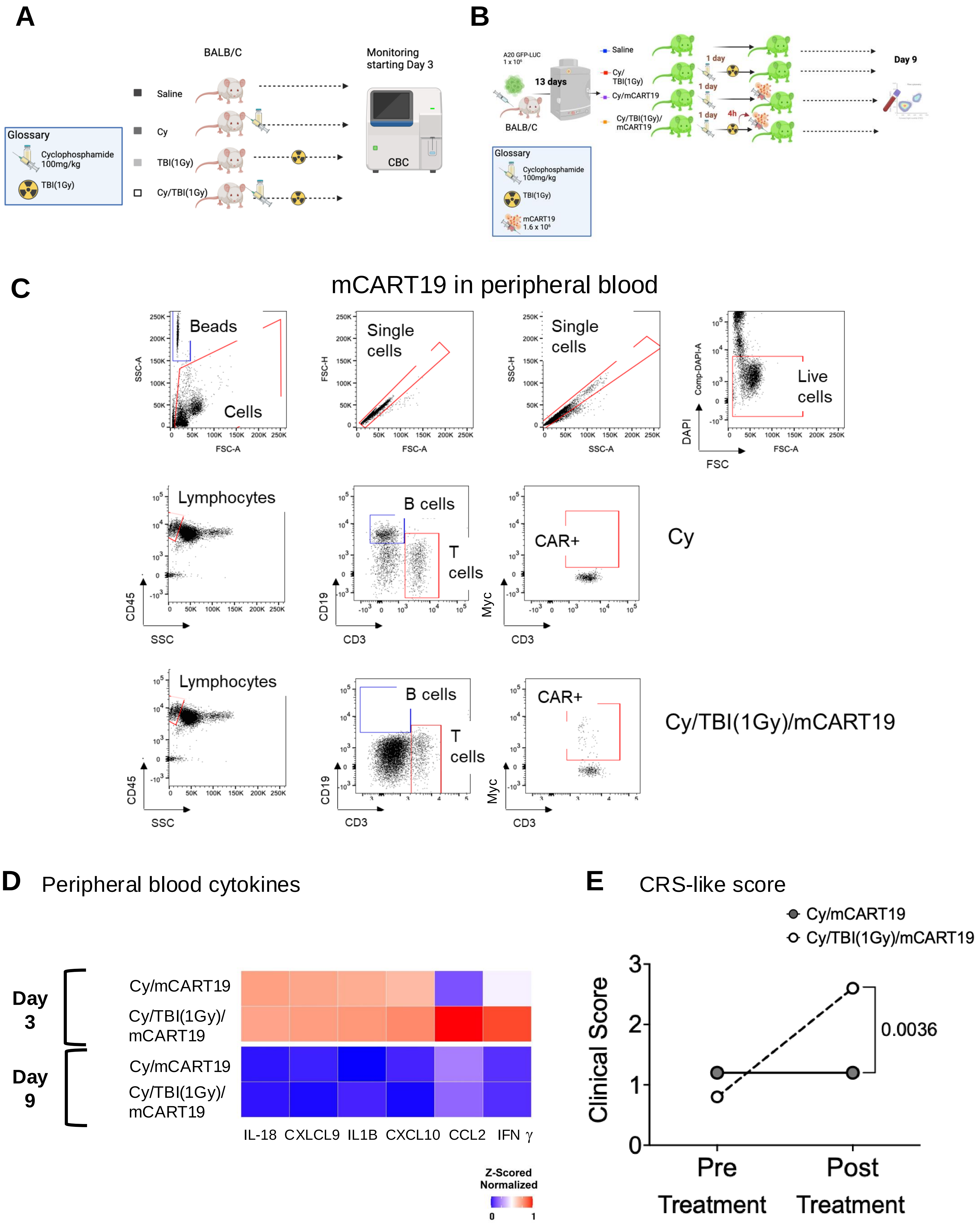

### supplementary Fig. 5

**Supplementary Figure 5.**

**A**

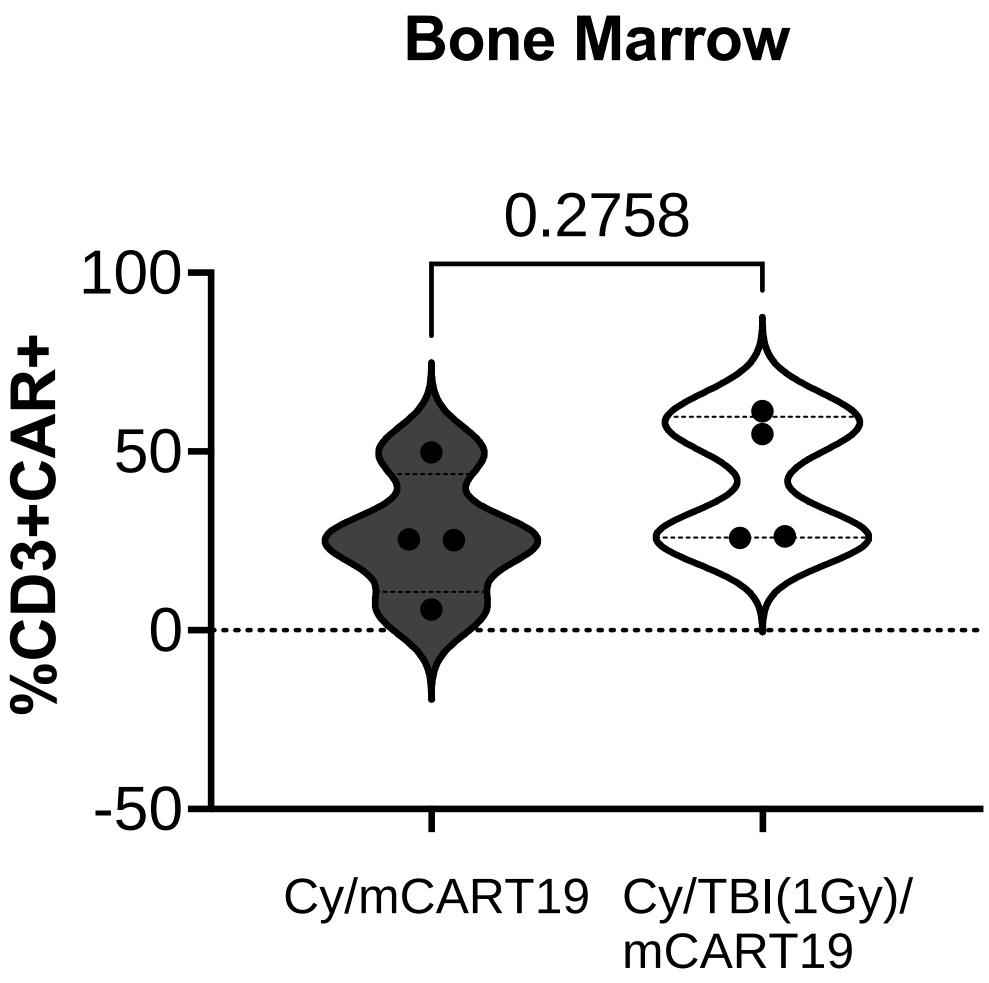


**B**

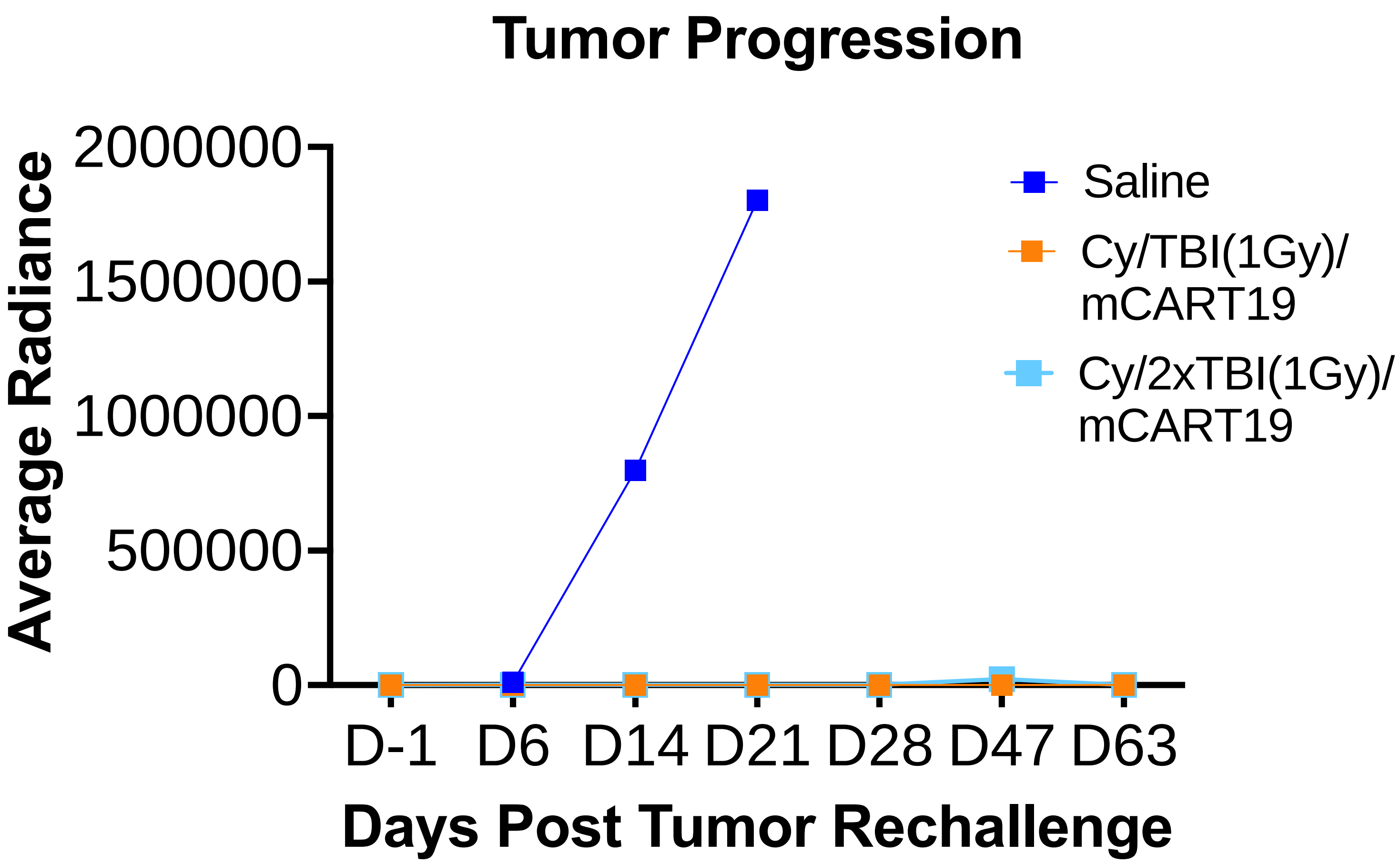

### supplementary Fig. 6A

#### Supplementary Figure 6 A.

**Naïve_1 T cell**
(Cy/TBI(1Gy)/mCART19 vs Cy/mCART19)

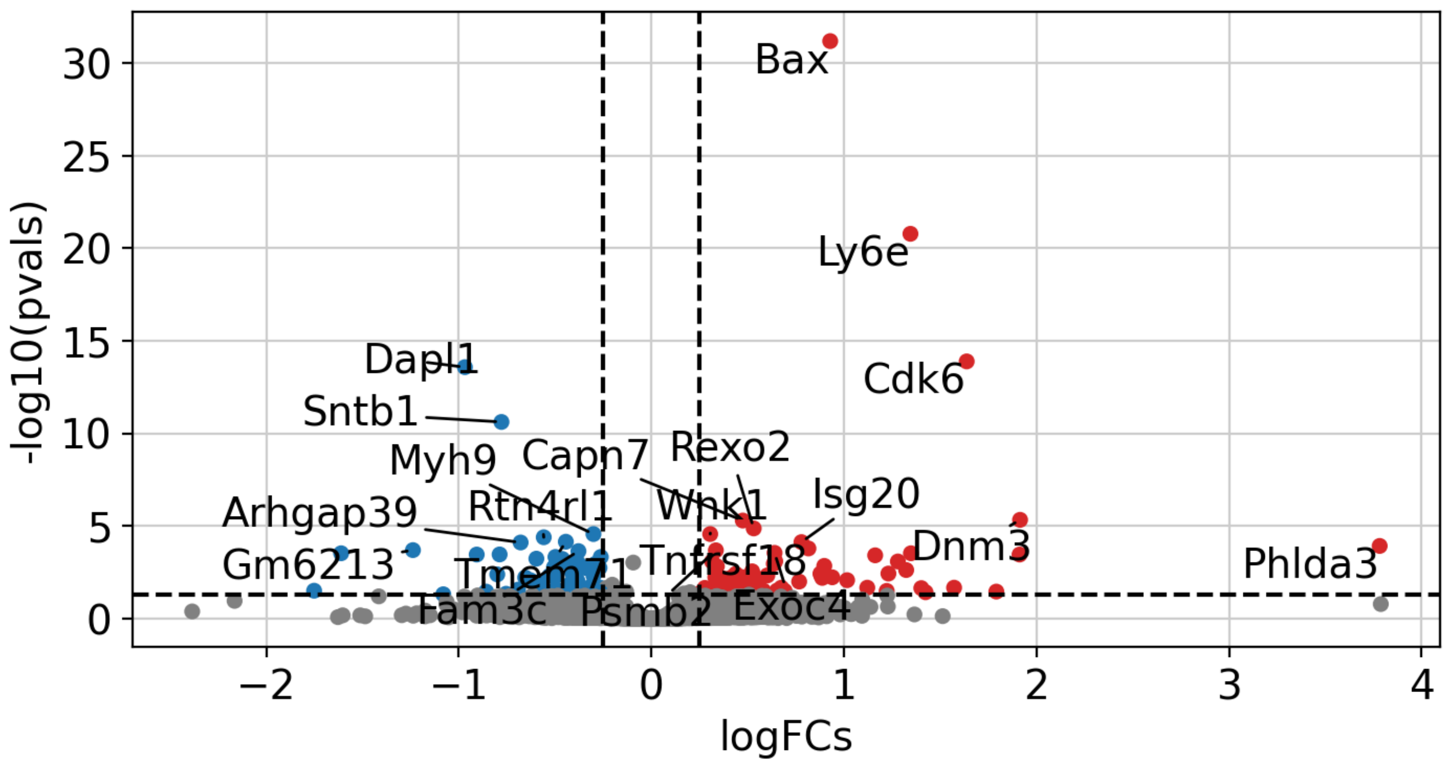


##### Enriched pathways- naïve_1 T cell

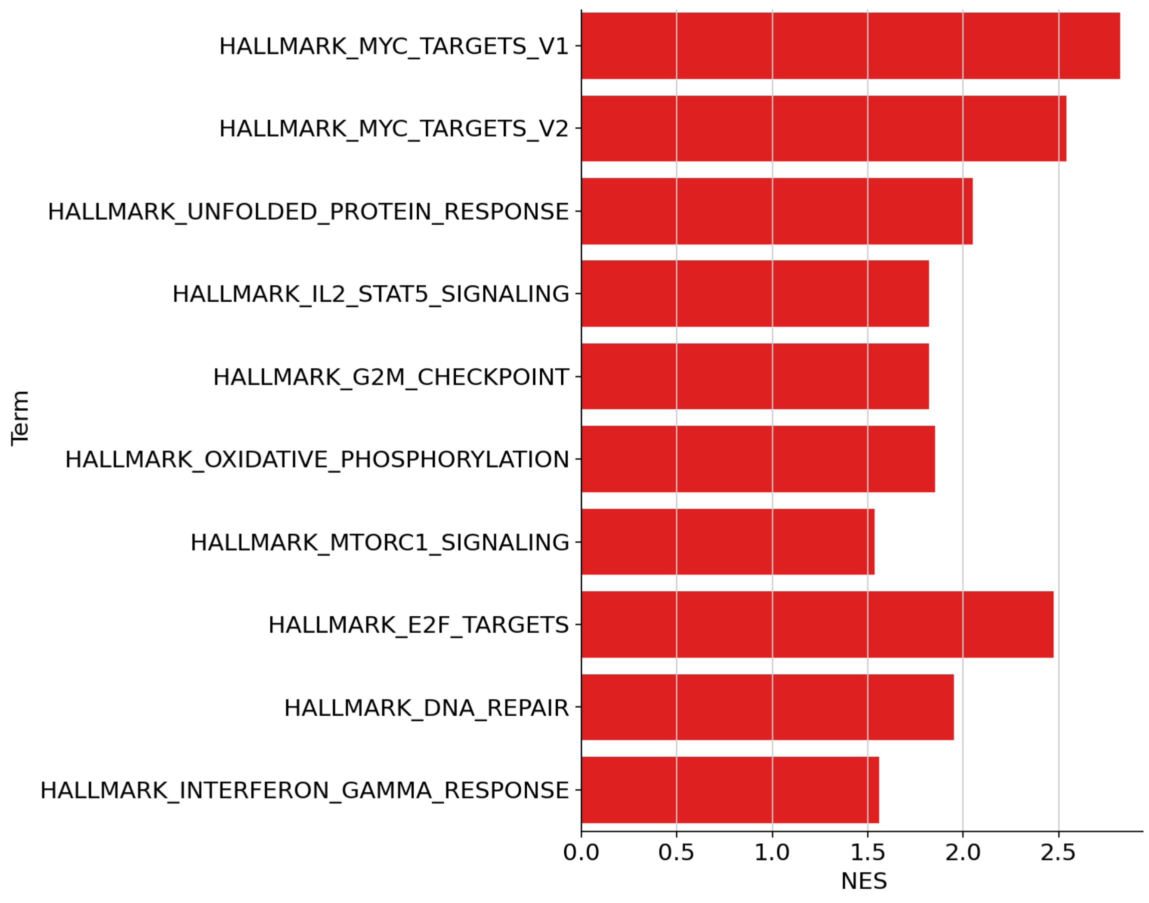

### supplementary Fig. 6B

**Supplementary Figure 6 B.**

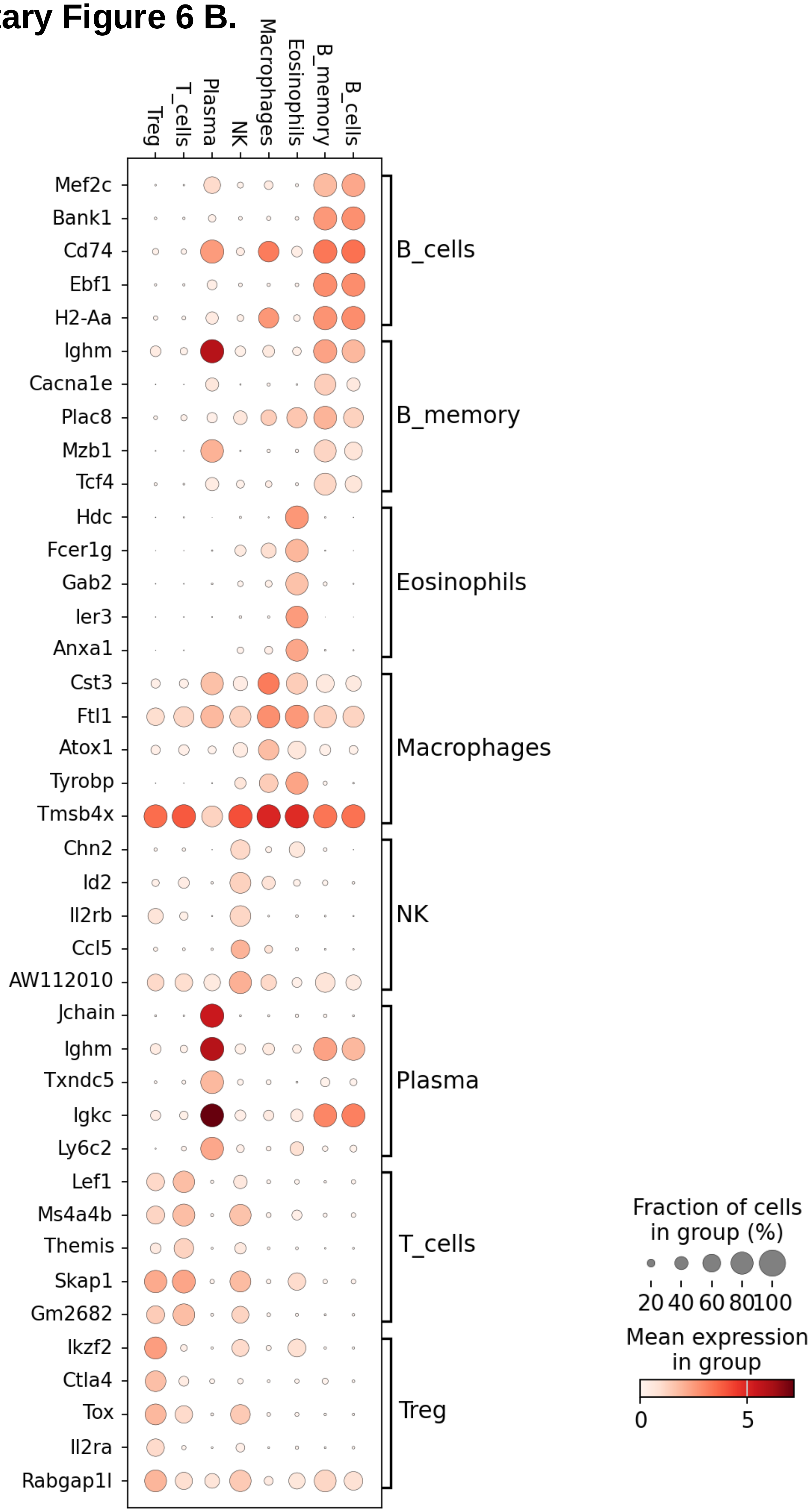

### supplementary Fig. 6C

#### Supplementary Figure 6 C.

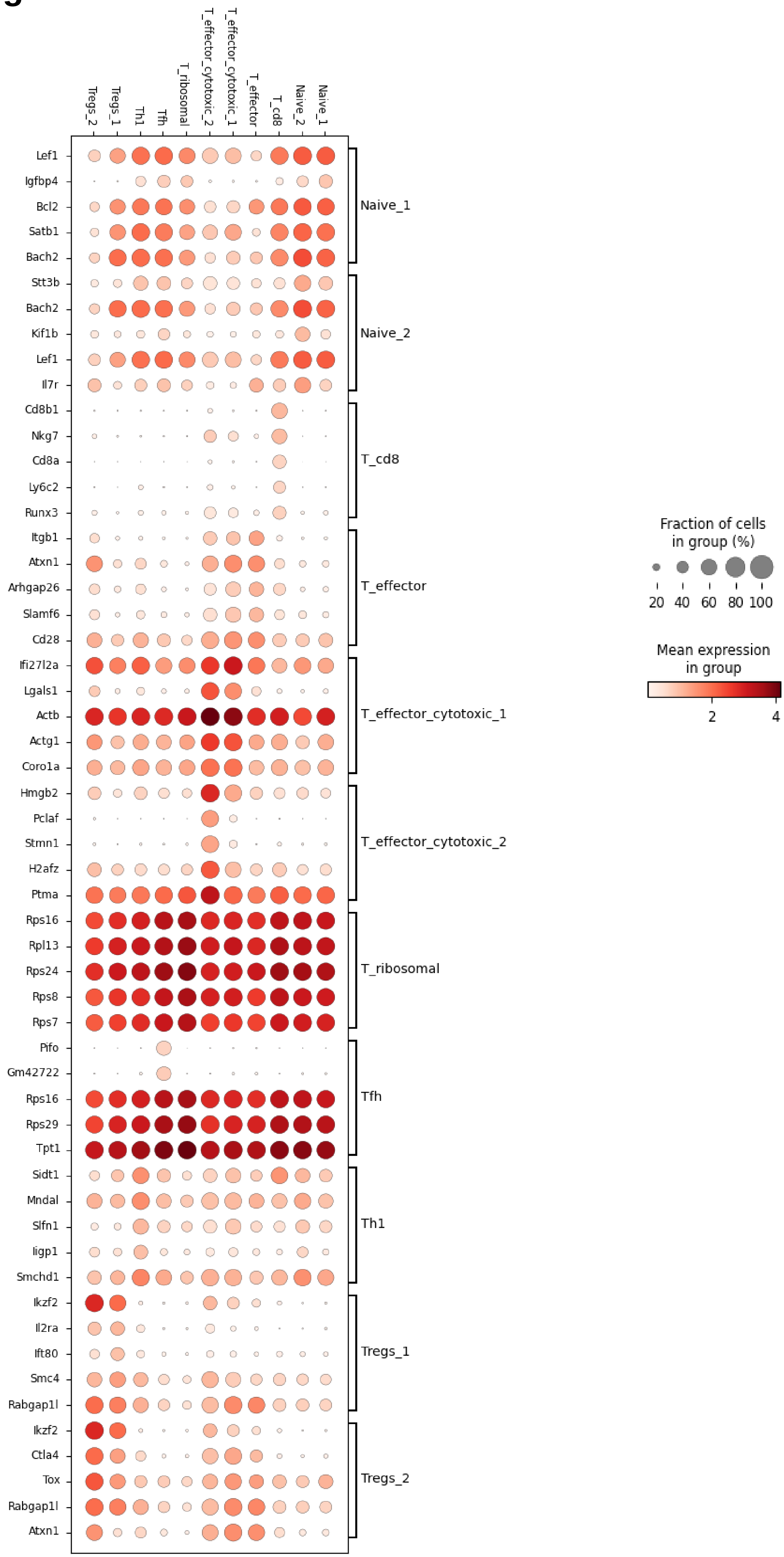
