## Supplementary Table 1. for "Total body irradiation primes CD19-directed CAR T cells against large B-cell lymphoma"

**Supplementary Table 1.** Cytokine release syndrome scoring system in mice

| Point | Hunched posture | Ruffled fur | >5% Weight loss |
| --- | --- | --- | --- |
| **0** | None | None | None |
| **1** | Present | Present | Present |
