## Supplementary Table 2. for "Total body irradiation primes CD19-directed CAR T cells against large B-cell lymphoma"

**Supplementary Table 2.** Flow cytometry antibodies and reagents

| Antibody | Vendor | Catalog number |
| --- | --- | --- |
| 4′,6-diamidino-2-phenylindole (DAPI) | Life Technologies | D3571 |
| CellTrace™ CFSE Cell Proliferation Kit | Life Technologies | C34554 |
| Anti-human TRAIL R2 BV510 clone B-K29 | BD OptiBuild | 745057 |
| Anti-human Fas Antibody clone ZB4 | Sigma | 05-338 |
| Anti-human FAS (CD95) PeCy7 clone DX2 | BD Pharmingen | 560177 |
| Anti-human CD19 PE clone SJ25C1 | BD Pharmingen | 560177 |
| Anti-human CD19 PE Vio 770 | Miltenyi | 130-113-647 |
| Anti-human CD45 Vio-blue | Miltenyi | 130-110-637 |
| Anti-human CD4 Vio-green | Miltenyi | 130-113-230 |
| Anti-human CD8 APC Vio 770 | Miltenyi | 130-110-681 |
| Anti-human CD3 FITC | Miltenyi | 130-113-138 |
| Anti-human EGFR PE (Hu1) | R&D | FAB9577P |
| Anti-mouse CD3 APC/Cy7 clone 17A2 | BioLegend | 100222 |
| Anti-mouse Myc PE clone 9B11 | Cell Signature | 3739S |
| Anti-mouse CD45 PE/Cy5, clone 30-F11 | BioLegend | 103110 |
| Anti-mouse CD8a APC clone 53-6.7 | BioLegend | 100712 |
| Anti-mouse CD19 BV650 clone 6D5 | BioLegend | 115541 |
| Anti-mouse CD4 PerCP-Cy5.5 clone RM4-5 | BioLegend | 100540 |
| Anti-mouse FAS (CD95) BV605 clone SA367H8 | BioLegend | 152612 |
| Anti-mouse CD45.2 APC clone 104 | BioLegend | 109814 |
| Anti-mouse CD8a BV510 clone 53-6.7 | BioLegend | 100751 |
| Anti-mouse CD19 PE Dazzle 594 clone 6D5 | BioLegend | 115554 |
| Anti-mouse CD44 PE-Cy5 clone IM7 | BioLegend | 103010 |
| Anti-mouse CD62L AF700 clone MEL-14 | BioLegend | 104426 |
| Anti-mouse IL7R- α (CD127) BV650 clone A7R34 | BioLegend | 135043 |
| Anti-mouse PD-1 CD279 PE-Cy7 clone 29F.1A12 | BioLegend | 135215 |
| YOPRO-1 iodine | Life Technologies | Y3603 |
